# Pooled AAV-CRISPR Screen with Targeted Amplicon Sequencing

**DOI:** 10.1101/153643

**Authors:** Guangchuan Wang, Ryan D. Chow, Lupeng Ye, Christopher D. Guzman, Xiaoyun Dai, Matthew B. Dong, Feng Zhang, Phillip A. Sharp, Randall J. Platt, Sidi Chen

**Author notes:** Co-first authors (Guangchuan Wang, Ryan D. Chow). Correspondence: SC, RJP.

## Abstract

High-resolution, high-throughput direct in vivo screening of functional genetic factors in native tissues has long been challenging. Adeno-associated viruses (AAV) are powerful carriers of transgenes and have been shown to mediate efficient genome editing in various organs in mice. Here, we developed a new technological approach, <u>P</u>ooled <u>A</u>AV-CRISPR <u>S</u>creen with <u>T</u>argeted <u>A</u>mplicon <u>S</u>equencing (PASTAS), and demonstrated its application for directly mapping functional cancer driver variants in the mouse liver in an autochthonous manner. Intravenous delivery of an AAV-CRISPR library targeting a set of the most frequently mutated tumor suppressor genes into fully immunocompetent conditional Cas9 knock-in mice consistently generated highly complex autochthonous liver tumors. The molecular landscapes of these genetically diverse tumors were mapped out by deep direct readout of Cas9-generated variants at predicted sgRNA cut sites using molecular inversion probe sequencing. Co-occurrence and correlation analyses as well as validation with lower complexity minipools further confirmed the potency of various co-mutated drivers. The PASTAS method can be applied to virtually any gene sets, any cancer types, or any type of in vivo genetic studies other than cancer.

## Introduction

Identification of causative variants in vivo is a key to understand genetic regulation of physiological or pathological processes ^1^. For example, linking phenotypes to driver genes is a central problem in cancer genetics ^2^. As the microenvironment is increasingly recognized to have a critical influence on tumorigenesis and progression ^3,4^, genetic screens or cancer models that rely on cell line transplants might not represent normal physiology ^5,6^. Autochthonous modeling of cancer – that is, in the native tissue of origin, provides greater translational relevance for cancer modeling and pre-clinical therapeutic studies. Genetically engineered mouse models (GEMMs) enables *in vivo* modeling of a wide variety of cancer types, and have become a cornerstone in studying the mechanisms of oncogenes and tumor suppressors *in vivo* ^5–7^. However, the production of GEMMs is time-consuming and requires a complex multi-step process ^8^, and GEMMs have largely been limited to the study of only a handful of genes at a time due to the technical difficulties of breeding large numbers of genetic modified mice.

Recent developments in genome editing utilizing CRISPR ^39–42^ enables modeling OGs and TSGs in somatic cells in various cancer types, which thus far have been demonstrated in only a few genes due to the challenges in generating autochthonous cancer models ^43,44,45^. Viruses have also been used to generate loss-of-function or gain-of-function mutations in tumor suppressors and oncogenes *in vivo* ^9,46–50^. However, these studies were limited to a small set of genes due to technological challenges and the nature of biological complexity in native organs. Thus far, CRISPR has been used to perform genome-scale knockout screens in cell lines and in transplant models ^51–54^, but not yet in the autochthonous setting of a native tissue. Moreover, current screens rely on the sequencing of sgRNAs, which is an indirect measurement of the selective forces acting on specific gene perturbations. Direct *in vivo* parallel mutational analysis of causative genetic variants generated by CRISPR has remained a challenge, necessitating the development of a new technological platform with higher throughput and precision.

To directly interrogate the comparative selective advantage of mutants in the tumor-initiating organs, it is necessary to first generate and then subsequently sequence pools of mutant cells within the native tumor environment. Adeno-associated viruses (AAV) are powerful carriers of transgenes and have been shown to mediate efficient genome editing in various organs in mice ^9,10^. Given that AAVs can efficiently infect the liver after intravenous injection ^11^, we reasoned that liver hepatocellular carcinoma (LIHC, also known as HCC), a deadly cancer with a poor five year survival rate ^12,13^, would be a suitable and relevant model for a proof-of-concept study.

Here, we developed a new technological approach, <u>P</u>ooled <u>A</u>AV-CRISPR <u>S</u>creen with <u>T</u>argeted <u>A</u>mplicon <u>S</u>equencing (PASTAS). We used this to directly map functional cancer genome variants of tumor suppressors in the autochthonous mouse liver using massively parallel CRISPR/Cas9 genome editing. We intravenously injected an AAV-CRISPR carrying a library of 278 sgRNAs that target a set of the most frequently mutated, known or putative tumor suppressor genes (TSGs) into *Rosa-LSL-Cas9-EGFP* knock-in mice (LSL-Cas9 mice) to generate highly complex autochthonous liver tumors, followed by direct readout of the Cas9-generated variants at predicted sgRNA cut sites using molecular inversion probe sequencing (MIPS). This combination of direct mutagenesis and pooled variant readout illuminated the mutational landscape of the tumors. Mutagenesis of individual or combinations of the top genes represented by high frequency variants led to liver tumorigenesis in fully immunocompetent mice, validating that PASTAS can be directly applied to quantify functional and causative genetic drivers.

## Results

To demonstrate the utility of the PASTAS method, we first develop it in a liver cancer model. We first sought to compile a list of the top SMGs in the pan-cancer TCGA datasets. Applying a similar approach as in previous studies ^14–16^, we identified the top 50 SMGs after excluding known oncogenes (Figure 1A). Of the top 50 putative TSGs, 49 genes had mouse orthologs (mouse TSGs, hereafter referred to as mTSG). We also selected seven housekeeping genes to serve as controls. Then, we designed a library of sgRNAs targeting these 56 different genes, with five sgRNAs for each gene, totaling 280 sgRNAs (hereafter referred to as the mTSG library) (Figure 1A; Table S1). For *Cdkn2a* and *Rpl22*, only four unique sgRNAs were synthesized, with the fifth sgRNA being a duplicate. The duplicates were treated as identical in downstream analyses. After oligo synthesis, we cloned the mTSG library into a base vector containing a U6 promoter driving the expression of the sgRNA cassette, as well as a Cre expression cassette (Figure 1A). Because mutation of a single TSG rarely leads to rapid tumorigenesis in humans or autochthonous mouse models, we included an sgRNA targeting *Trp53* in the base vector, with the initial hypothesis that concomitant *Trp53* loss-of-function might facilitate tumorigenesis. Sequencing of the plasmid pool revealed a complete coverage of the 278 unique sgRNAs represented in the mTSG library (Table S2). After generating AAVs (serotype AAV9) containing the base vector or the mTSG library, we then intravenously injected PBS, vector AAVs or mTSG AAVs into fully immunocompetent LSL-Cas9 mice (Figure 1A). Upon AAV infection, Cre is expressed and excises the stop codon, activating Cas9 and EGFP expression.

**Figure 1:**
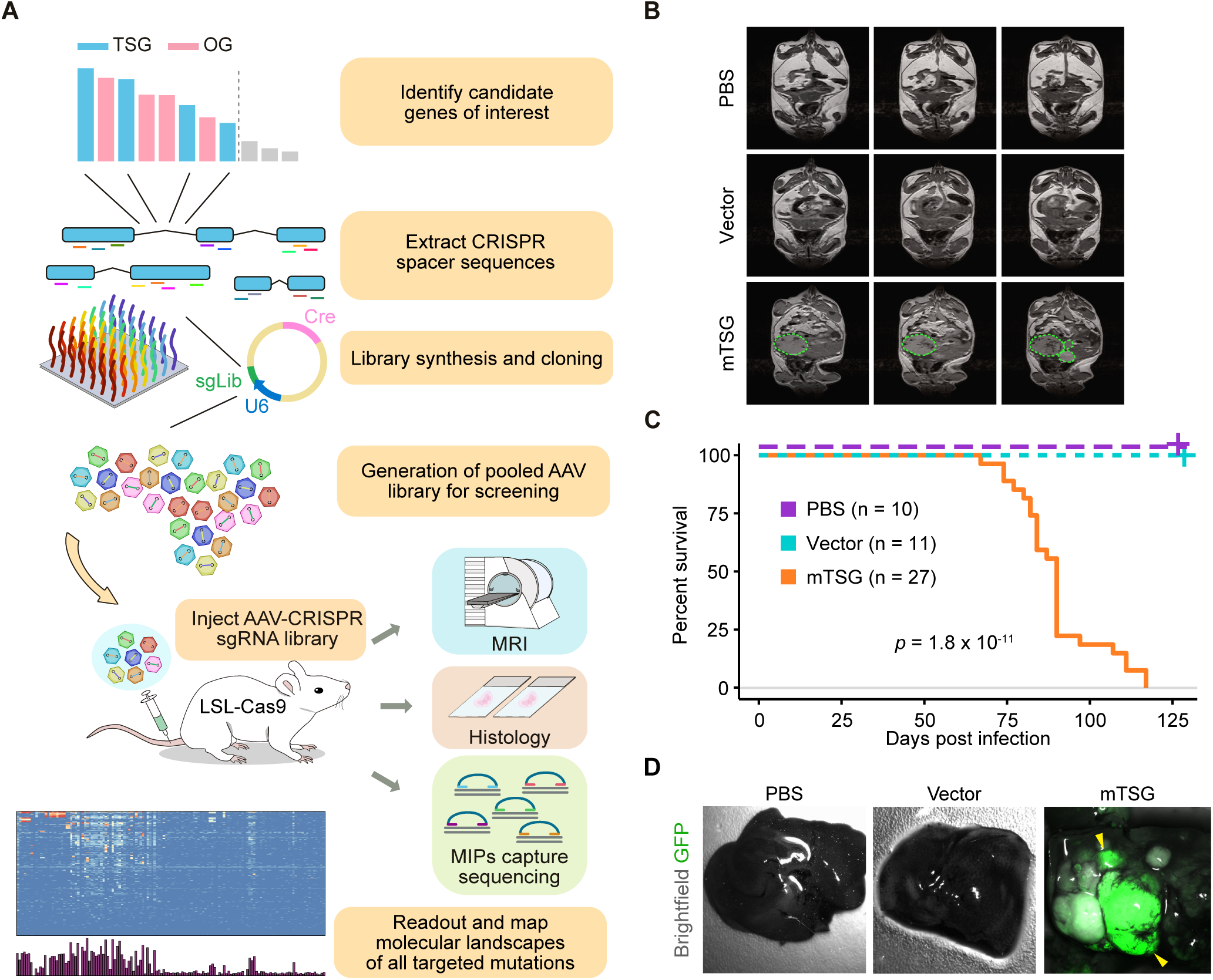
Development and demonstration of Pooled AAV screen with Targeted Amplicon Sequencing (PASTAS) in an autochthonous liver cancer model. **A.** Schematics of the overall design and experimental outline of PASTAS in autochthonous LIHC study. First, the top significantly mutated genes were identified from pan-cancer TCGA datasets. After removing known oncogenes and genes without mouse orthologs, a set of 49 most significantly mutated putative tumor suppressor genes were chosen (mTSG). Seven additional genes with house-keeping functions were spiked-in, leading to a final set of 56 genes. SgRNAs targeting these genes were then identified computationally and five were chosen for each gene. 280 sgRNAs plus eight non-targeting control (NTC) sgRNAs were synthesized, and then the sgRNA library (mTSG, 288 sgRNAs) was cloned into an expression vector that also contained Cre recombinase and a *Trp53* sgRNA. AAVs carrying the mTSG library were produced and injected into the tail veins of LSL-Cas9 mice. After a specified time period, the mice were subjected to Magnetic resonance imaging (MRI), histology, and MIPs capture sequencing for readout and deep variant analysis of molecular landscape of all targeted genes and mutations. **B.** MRI of abdomens of mice treated with PBS, vector, or mTSG library. Detectable tumors are circled with green dashed lines. PBS treated mice (n = 3) did not have any detectable tumors, while vector treated mice (n = 3) occasionally had small nodules. In contrast, mTSG-treated mice (n = 4) often had multiple detectable tumors. **C.** Kaplan-Meier survival curves for PBS (purple, n = 10), vector (teal, n = 11), and mTSG (orange, n = 27) treated mice. No mTSG-treated mice survived longer than four months post treatment, while all PBS and vector treated animals survived the duration of the experiment. Statistical significance was assessed by log-rank test (*p* = 1.8 * 10^-11^). **D.** Brightfield images with GFP fluorescence overlay (green) of livers from representative PBS, vector, and mTSG-treated mice, 4 months post-treatment. Large GFP+ tumors are marked with yellow arrowheads. In contrast to PBS or vector-treated mice, mTSG-treated mice had numerous detectable GFP+ nodules.

Live magnetic resonance imaging (MRI) of mice 3 months post-treatment revealed large nodules in mTSG-treated animals (n = 4), while vector-treated animals (n = 3) only occasionally had small nodules and PBS animals (n = 3) were devoid of detectable nodules (Figure 1B; Figure S1A-B; Table S3). The total tumor volume in each mouse was significantly larger in mTSG samples compared to PBS and vector samples (one-sided Mann-Whitney test, *p* = 0.0286 and *p* = 0.0286, respectively) (Figure S1B). These data suggest that the AAV-CRISPR mTSG library is sufficient to induce rapid tumorigenesis in the livers of LSL-Cas9 transgenic mice.

Mice that received the AAV-CRISPR mTSG library (n = 27) did not survive more than four months (median survival = 90 days; 95% confidence interval = 84 - 90 days), while mice that were treated with PBS (n = 10) or vector control (n = 11) all survived the duration of the experiment (log-rank test, *p* = 1.8 * 10^-11^) (Figure 1C; Table S4). By gross examination under a fluorescent dissecting scope, detectable GFP+ nodules were observed in mTSG-treated livers, but not in PBS or vector samples (Figure 1D; Figure S2). Notably, in mTSG-treated mice, we occasionally observed tumors that were not primarily located in the liver. Chief among these were several big abdominal tumors (BATs, n = 6), as well as a few sarcomas (n = 4) and ear tumors (n = 2), although BATs were later found to be of liver origin on the basis of histological analysis.

We analyzed endpoint histological sections from PBS (n = 7), vector (n = 5), and mTSG-treated mice (n = 13), sacrificed 3-4 months post-treatment (Figure 2A; Figure S3-4). No tumors were found in PBS-treated mice, while rare small tumors were found in vector-treated mice (total tumor area = 5.96 ± 3.27 mm^2^) (Fig. 2B). Consistent with the MRI results, mice that received the mTSG library had significantly larger liver tumors, with the pathology of LIHC (total tumor area = 100.6 ± 47.19 mm^2^; one-sided Welch’s t-test, *p* = 0.027 compared to PBS, *p* = 0.034 compared to vector) (Figure 2A-B; Table S5). As these mice were found to have multiple liver tumors, we also compared the size of each individual tumor across the 3 treatment groups (Figure 2C). The mTSG-treated mice collectively had tumors that were significantly larger (26.69 ± 6.18 mm^2^) than the tumors found in vector-treated animals (3.31 ± 1.55 mm^2^; *p* = 0.0003), though the latter were too small to be detected by gross examination under GFP dissecting scope. We assessed the proliferation of liver samples from PBS, vector, and mTSG-treated mice by Ki67 expression, and found that rapid proliferation was restricted to tumor cells (Figure S4B). Additionally, we found that the tumors in mTSG treated mice, but not vector treated mice, were largely positive for AE1/AE3 (pancytokeratin), which is a marker of LIHC (Figure 2D; Figure S4C). These data collectively indicate that the AAVCRISPR mTSG library directly promotes aggressive liver tumorigenesis in otherwise wildtype LSL-Cas9 mice.

**Figure 2:**
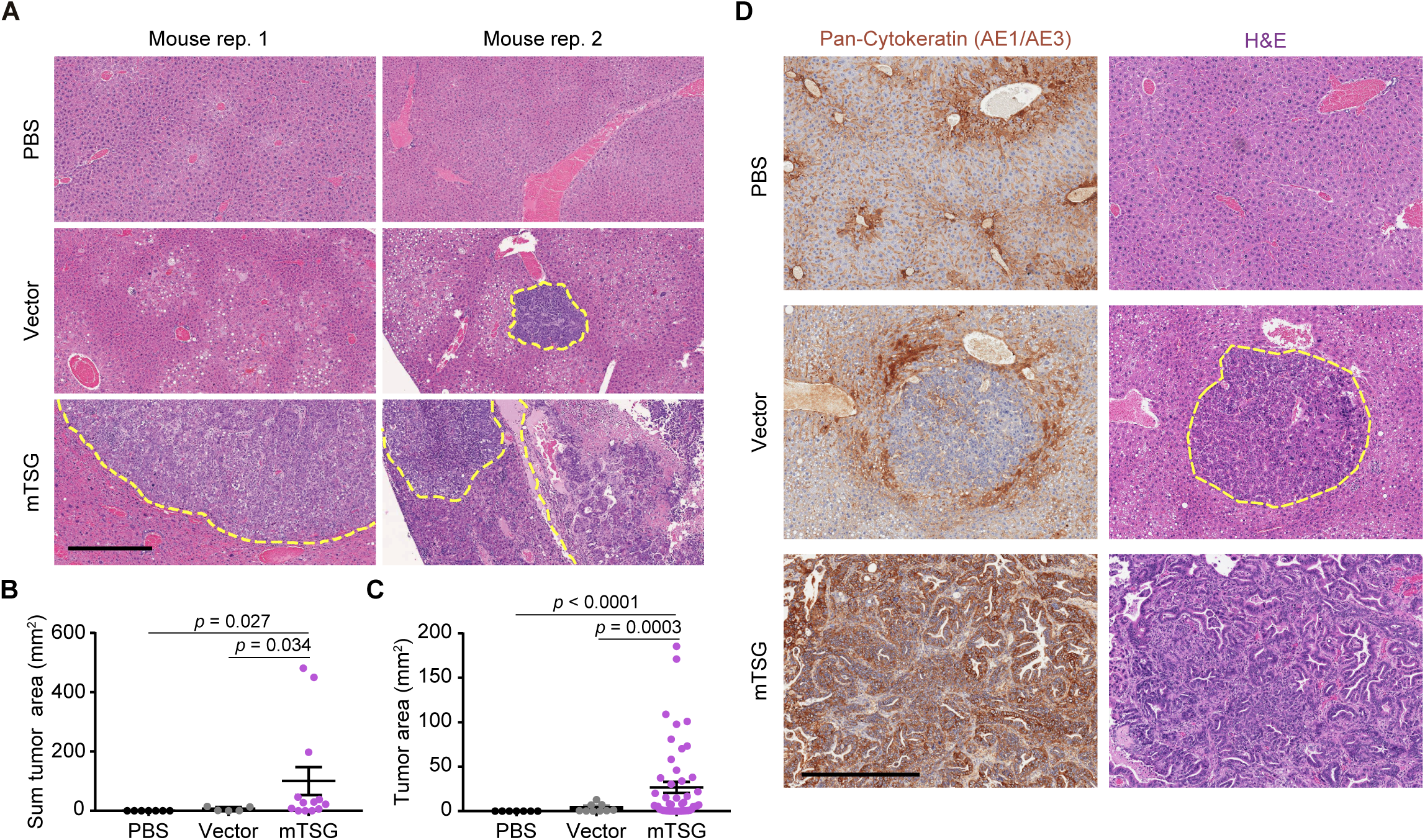
Histology analysis of autochthonous tumors generated by AAV-CRISPR mTSG library. **A.** Hematoxylin and eosin staining of liver sections from mice treated with PBS (n = 7), vector (n = 5), or mTSG library (n = 13). Tumor-normal boundaries are demarcated with yellow dashed lines. No tumors were found in PBS samples, while small nodules were found, although rare, in vector samples. On the other hand, mTSG-treated livers were replete with tumors (statistics in **B-C**). **B.** Dot plot of the total tumor area per mouse (mm^2^) in liver sections from mice treated with PBS (black, n = 7), vector (gray, n = 5), or mTSG library (purple, n = 13). mTSG-treated mice had a significantly higher total tumor burden than PBS (one-sided Welch’s t-test, *p* = 0.027) or vector-treated mice (*p* = 0.034). **C.** Dot plot of the individual tumor area (mm^2^) in liver sections from mice treated with PBS (black, n = 7), vector (gray, n = 9), or mTSG library (purple, n= 49). mTSG-treated mice had significantly larger tumors than PBS (one-sided Welch’s t-test, *p* < 0.0001) or vector-treated mice (*p* = 0.0003). **D.** Representative IHC staining of a LIHC marker, pan-cytokeratin (AE1/AE3) from mice treated with PBS, vector, or mTSG library. The tumors from mTSG-treated samples shown revealed positive staining for AE1/AE3, consistent with LIHC pathology. Certain mTSG tumors were partially positive for cytokeratin, revealing tumor heterogeneity. The tumors from vector-treated samples were relatively small and almost always negative or slightly positive for cytokeratin. Scale bar is 0.5 mm.

To understand the molecular alterations driving the development of tumors in mTSG-treated mice, we designed molecular inversion probes (MIPs) to enable capture sequencing of the ±70 basepair (bp) regions surrounding the predicted cut site of each sgRNA in the mTSG library (namely, the +17 position of each 20 bp spacer sequence) (Methods). As opposed to simply sequencing the sgRNA cassettes to find the relative enrichment of each sgRNA within the cell population, MIP capture-sequencing enables a direct quantitative analysis of the mutations induced by the Cas9-sgRNA complex. To generate this pool of MIPs (termed mTSG-MIPs) (Table S6), we synthesized a total of 266 extension and ligation probes targeting 266 genomic loci with an average size of 158 ± 8 (SEM) bp, covering 278 unique sgRNA sites. Liver genomic DNA was extracted from PBS-treated (n = 8 mice), vector-treated (n = 8 mice), and mTSG-treated mice (n = 27 mice; 37 liver lobes in total). In order to assess the potential for AAV-CRISPR mediated mutagenesis of other organs, we also collected DNA from all observed non-liver tumors (n = 23), as well as a wide variety of tissues (such as brain, lung, colon, spleen and kidney) without detectable tumors under a fluorescent dissecting scope (n = 57 samples) from all three groups. We performed MIP capture-sequencing on all genomic DNA samples (total n = 133; Table S7). Sequencing depth of the sgRNA target regions was sufficiently powerful to detect variants at < 0.01% frequency, with a mean read depth of 13,286 ± 1033 (SEM) across all MIPs after mapping to the mouse genome (Table S8). Median read depth across all MIPs approximated a log-normal distribution, indicating relatively even capture of the target loci (Figure S5a). Insertions and deletions (indels) were then called across all samples to reveal detectable indel variants at each sgRNA cut site (Methods) (Table S9). We excluded single nucleotide variants (SNVs) from the analysis, as indels are the dominant variants generated by non-homologous end-joining (NHEJ) following Cas9-mediated double-strand breaks (DSBs) *in vivo* ^17–19^. For downstream analysis, we only considered indels that overlapped the ±3 bp flanks around each of the predicted sgRNA cut sites, as Cas9 tends to create DSBs within a tight window near the predicted sgRNA cut site in mammalian cells ^19^. A representative example of the genotypes observed by MIPs capture-sequencing is shown at the *Setd2* sgRNA 1 cut site for PBS, vector or mTSG-treated samples (Figure 3A), illustrating the diversity of Cas9-induced indels in mTSG-treated mice.

**Figure 3:**
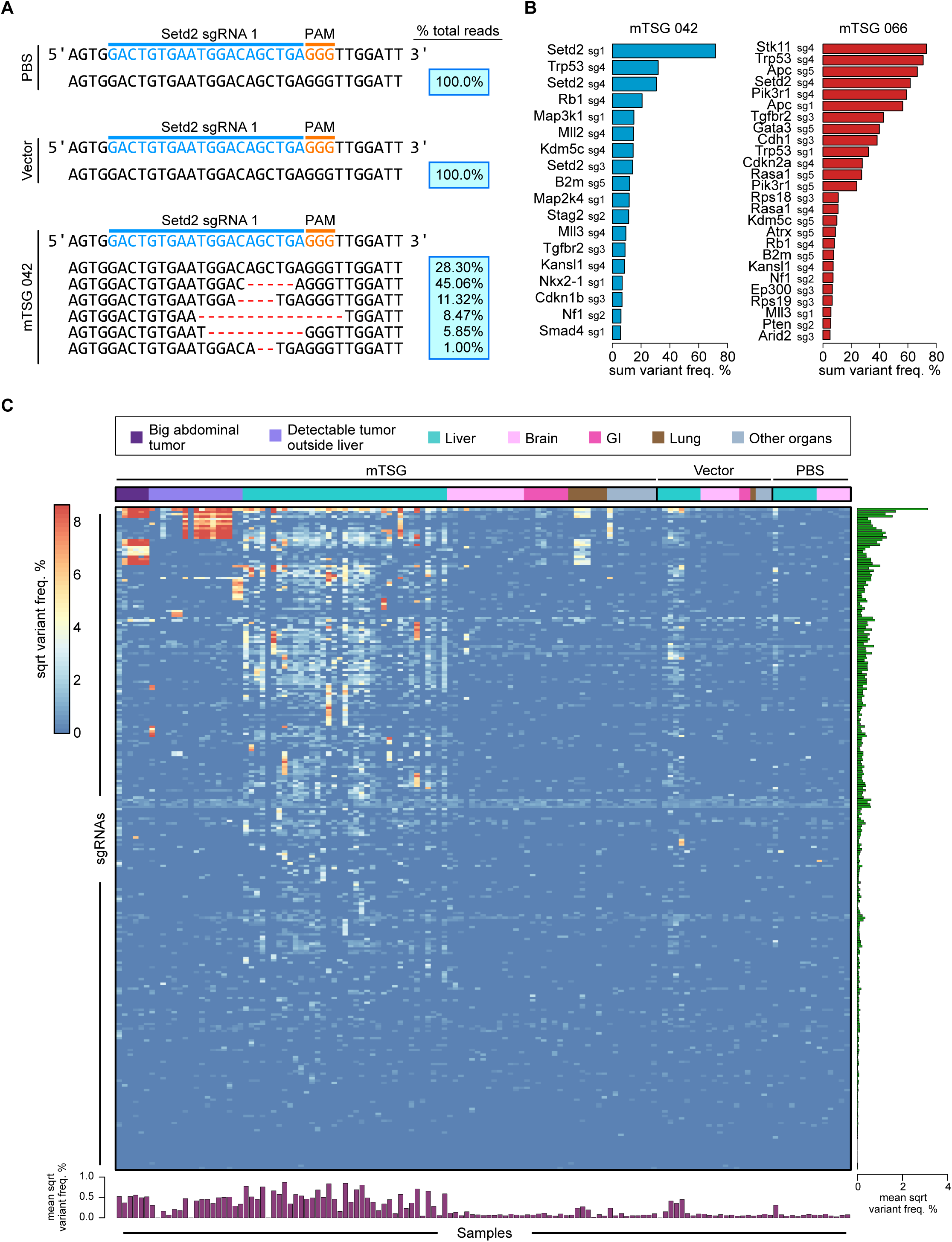
Mutational variant level mutational landscape of LIHC generated by PASTAS technology. **A**. Unique variants observed at the genomic region targeted by *Setd2* sg1 in representative PBS, vector, and mTSGtreated liver samples. The percentage of total reads that correspond to each genotype is indicated on the right in the blue boxes. No indels were found in the PBS or vector-treated samples, while several unique variants were identified in the mTSG-treated sample (mTSG 042). **B.** Waterfall plots of two mTSG-treated samples (042, 066) detailing sum variant frequencies in significantly mutated sgRNA sites (SMSs). Individual mice presented with distinct mutational signatures, suggesting that a wide variety of mutations induced by the mTSG library had undergone positive selection. **C.** Global heatmap detailing the square-root of sum variant frequency across all sequenced samples (n = 133) from mTSG (n = 98 samples), vector (n = 21 samples) or PBS-treated mice (n = 14 samples) in terms of sgRNAs. Square-root transformation was used to even out the distribution of variant frequencies for visualization. Each row represents one sgRNA, while each column represents one sample. Treatment conditions and tissue type are annotated at the top of the heatmap: big abdominal tumor (dark purple), detectable tumor outside liver (light purple), liver (teal), brain (light pink), gastrointestinal (GI) (dark pink), lung (brown) and other organs (gray). Bar plots of the mean square-root variant frequencies for each sgRNA (right panel, green bars) and each sample (bottom panel, purple bars) are also shown. mTSG-treated organs without visible tumors (0.148 ± 0.037 SEM) had significantly lower mean variant frequencies compared to mTSG-treated tumors and livers (BATs, 3.098 ± 0.600; two-sided unpaired t-test, *p <* 0.0001), non-liver tumors (1.919 ± 0.338; *p <* 0.0001), and livers (1.451 ± 0.203; *p <* 0.0001). Livers and other organs from vector-treated animals (0.398 ± 0.179 and 0.054 ± 0.004, respectively) and PBS-treated animals (0.140 ± 0.067 and 0.063 ± 0.021, respectively) all had significantly lower variant frequencies than mTSG-treated livers (*p* < 0.0001 for all comparisons).

After collapsing each of the filtered indel calls to the closest sgRNA by summing their constituent variant frequencies (Tables S10), we plotted the overall spectrum of variant frequencies across all sequenced samples (Figure 3C). We then calculated the mean variant frequency for each sgRNA and each sample (right and bottom panels, respectively) (Figure 3C). The mTSG-treated organs without visible tumors (0.148 ± 0.037 SEM) had significantly lower mean variant frequencies compared to mTSG-treated tumors and livers (BATs, 3.098 ± 0.600; unpaired t-test, *p <* 0.0001), non-liver tumors (1.919 ± 0.338;*p <* 0.0001) and livers (1.451 ± 0.203;*p <* 0.0001). Livers and other organs from vector-treated animals (0.398 ± 0.179 and 0.054 ± 0.004, respectively) and PBS-treated animals (0.140 ± 0.067 and 0.063 ± 0.021, respectively) all had significantly lower variant frequencies than mTSG-treated livers (*p* < 0.0001 for all comparisons). The low background variant frequencies observed in vector-treated and PBS-treated samples may be due to noise that was generated during sequencing, as well as stochastic or germline mutations. Of note, the vector contains a *Trp53* sgRNA that potentially contributes to higher variant frequencies in vector-treated livers due to genome instability of *Trp53*-deficient cells.

We identified significantly mutated sgRNA sites (SMSs) in the mTSG-treated liver samples using a false-discovery rate (FDR) method as compared to PBS and vector-treated liver samples, such that no control sample would have any called SMSs (Methods). As we were most interested in analyzing dominant clones that had undergone strong positive selection in the tumor, we further required that at least 5% of the reads must have an indel in that region in order to call an SMS (Table S11). Interestingly, different mTSG-treated liver samples presented with highly heterogeneous mutational signatures, indicating that a diverse array of mutations had undergone positive selection in different samples (Figure S6).

We then collapsed SMSs in each sample to gene level to find significantly mutated genes (SMGs) (Table S12). Analysis of all mTSG liver samples revealed a full mutational landscape of the entire cohort (Figure 4, Figure S7). Out of 37 mTSG-treated liver samples, 33 (89%) were found to have major indels (≥ 5% sum variant frequency and FDR < 0.0625) (Methods) in one or more of the 56 genes in the mTSG library (average number of SMGs per sample = 11.7 ± 1.53). *Trp53*, *Setd2*, *Cic*, and *Pik3r1* were the top mutated genes in the cohort (mutated in 24/37, 18/37, 17/37 and 17/37 samples, respectively). *Trp53* is a well-known tumor suppressor that has been found to directly induce liver tumors upon loss-of-function in hepatocytes ^20^; *Setd2* is an epigenetic modifier that has been implicated in clear cell renal carcinoma ^21^, but not yet functionally characterized in liver cancer; *Cic* is a transcriptional repressor that has been shown to be a negative regulator of EGFR signaling ^22–24^; *Pik3r1* is a modulator of PI3K signaling, and loss-of-function mutations of this gene have been found to induce liver tumorigenesis in mice ^25^. In terms of cellular pathways, epigenetic modifiers and cell death/cell cycle regulators were frequently mutated, with multiple genes that were significantly mutated in more than 20% of samples (Figure 4). While the importance of epigenetic modifiers in cancer is now accepted ^26^, direct functional testing of groups of epigenetic regulators in an autochthonous model of tumorigenesis has not yet been shown in a systems manner.

**Figure 4:**
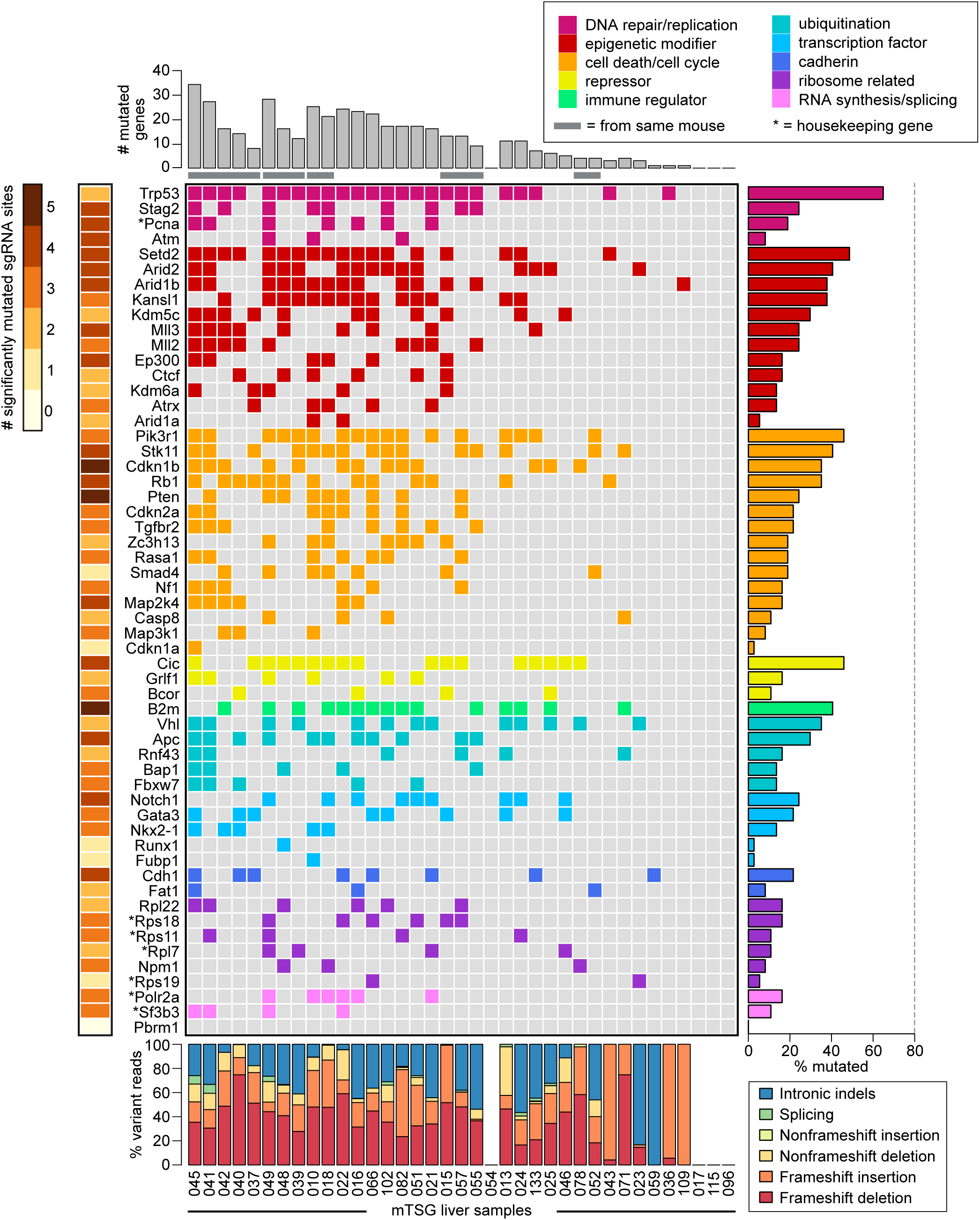
Mouse gene-level mutational landscape of LIHC generated by PASTAS technology. Each row in the figure corresponds to one gene in the mTSG library, while each column corresponds to one mTSGtreated liver sample. **Top:** Bar plots of the total number of significantly mutated genes (SMGs) identified in each mTSG-treated liver sample (n = 37). Samples originating from the same mouse are grouped together and denoted with a gray bar underneath. **Center:** Tile chart depicting the mutational landscape of primary liver samples infected with the mTSG library. Genes are grouped and colored according to their functional classifications (DNA repair/replication, epigenetic modifier, cell death/cycle, repressor, immune regulator, ubiquitination, transcription factor, cadherin, ribosome related and RNA synthesis/splicing), as noted in the legend in the top-right corner. Colored boxes indicate that the gene was significantly mutated in a given sample, while a gray box indicates no significant mutation. Asterisks denote several pre-selected genes that were generally considered housekeeping genes. **Right:** Bar plots of the percentage of liver samples that had a mutation in each of the genes in the mTSG library. *Trp53*, *Setd2*, *Pik3r1*, *Cic, B2m, Vhl, Notch1, Cdh1, Rpl22* and *Polr2a* were the top mutated genes in each of the 10 functional classifications, respectively. **Bottom:** Stacked bar plots describing the type of indels observed in each sample, color-coded according to the legend in the bottom-right corner. Frameshift insertions or deletions comprised the majority of variant reads (median = 59.2% across all samples). **Left:** Heatmap of the number of significantly mutated sgRNA sites (0-5 SMSs) for each gene. Multiple significantly mutated sgRNA sites for a given gene are indicative of a strong selective force for loss-of-function mutations in that gene.

Of the genes that were significantly mutated in at least one sample, the vast majority (91%, or 50/55) had multiple SMSs (median = 3 SMSs out of 5 total sgRNAs per gene), suggesting that these genes are indeed functional tumor suppressors (Figure 4). ANNOVAR analysis of the indels present in the mTSG liver cohort revealed that frameshift insertions and frameshift deletions comprised the majority of total variant reads (median = 59.2% across all samples) (Figure 4; Figure S5B), consistent with the notion that frameshift mutations are expected to cause loss-of-function in genes. Intronic, splice site and non-frameshift mutations nevertheless comprised a sizeable proportion of total variant reads (Figure 4). Thus, the PASTAS method can induce robust phenotypes and map out a molecular landscape of all targeted genes and genetic variants in an unbiased manner.

To explore synergistic effects between different genes in the mTSG library, we performed co-mutation analysis. For each pair of genes, by tabulating the number of samples that were double mutant, single mutant or double wildtype, we determined the strength of mutational co-occurrence (Methods) (Figure S8A). Out of all 1540 possible gene pairs, we found that a total of 226 pairs were significantly enriched beyond what would be expected by chance (hypergeometric test, *p* < 0.05), with highly significant pairs such as *Cdkn2a* + *Pten* (co-occurrence rate = 7/10 = 70%; hypergeometric test, *p* = 2.63 * 10^-5^), *Cdkn2a* + *Rasa1* (co-occurrence rate = 6/9 = 67%; *p* = 7.96 * 10^-5^), *Arid2* + *Cdkn1b* (co-occurrence rate = 11/17 = 65%; *p* = 9.13 * 10^-5^), and *B2m + Kansl1* (co-occurrence rate = 11/18 = 61%; *p* = 3.6 * 10^-4^) (Figure S8B-C; Table S13). Loss-of-function mutations in both genes of these combinations might synergistically enhance tumor progression.

We then investigated whether genes correlated with each other in terms of mutation frequencies within individual tumors. Since the variant frequency is essentially a metric for the positive selection that acts on a given mutation, genes whose variant frequencies are highly correlated across samples could also be synergistic in driving tumorigenesis. We calculated the total variant frequency for each gene by summing all the values from all five sgRNAs, used these summed gene level variant frequencies across each sample to calculate the Spearman correlation between all 1540 possible gene pairs, and assessed whether the correlations were statistically significant (Methods) (Figure S8D; Table S14). A total of 128 gene pairs were significantly correlated (Spearman correlation, Benjamini-Hochberg adjusted *p* < 0.05). The top four correlated pairs were *Cdkn2a* + *Pten* (Spearman R = 0.817, *p* = 6.97* 10^-10^), *Nf1* + *Rasa1* (R = 0.791, *p* = 5.86 * 10^-9^), *Arid2* + *Cdkn1b* (R = 0.788, *p* = 7.16 * 10^-9^), and *Cdkn2a* + *Rasa1* (R = 0.761, *p* = 4.45 * 10^-8^) (Figure S8E-F). We performed the same analysis using Pearson correlation, finding extensive similarities in the identified pairs (Figure S9A-B; Table S14). As the base vector contained a *Trp53* sgRNA, we also performed the co-mutation analyses excluding all pairs involving *Trp53* (Figure S9C-D). The correlation analysis thus revealed a number of highly significant associations in specific pairs of genes. Four gene pairs were statistically significant at Benjamini-Hochberg adjusted *p <* 0.05 in both the co-occurrence and correlation analyses (Figure S8G). Interestingly, one of the top gene pairs was *Arid2* + *Cdkn1b*, representing a previously unreported synergistic interaction between an epigenetic regulator and a cell cycle regulator.

To deepen our understanding of the clonal architecture in this genetically complex, highly heterogeneous yet fully gene-targeted autochthonous tumor model, we reframed our analysis to the level of specific indel variants. We focused on single mouse that had presented with multiple visible tumors in several liver lobes, five of which had been harvested for MIPs capture sequencing (Methods) (Figure 5A). Analysis of the sgRNA-level variant frequencies in the five lobes revealed strong pairwise correlations between multiple lobes (Figure 5B). Across all 37 mTSG-treated liver samples, we identified 593 unique variants that had a variant frequency ≥ 1% in at least one sample (Table S15). The majority of these variants (80.94%) were deletions rather than insertions (Table S15). The inter-lobe correlations are suggestive of similar variant compositions within these liver lobes. Hierarchical clustering of the variant-level data across all mTSG-treated liver samples revealed the existence of sample-specific variants. 70.15% (416/593) of the variants were sample-specific (private variants), while 29.85% (177/593) variants were found across multiple samples (shared variants) (Figure 5C). Shared variants could originate from convergent processes of NHEJ following Cas9/sgRNA mediated DSBs, leading to the same indel pattern, or alternatively from clonal expansion or metastasis. In an example mouse with five lobes MIPS-sequenced, we identified 178 unique variants (> 1% variant frequency threshold) represented within the five liver lobes (Table S16). Using binary variant calls (i.e., whether a given variant is present or absent in a sample), we clustered these 178 variants into eight groups (Figure 6D, Table S16). Variants in clusters 1, 2, 3, 5, and 6 were specific to a single lobe (private variant clusters), whereas variants in clusters 4, 7, and 8 were present across multiple lobes (shared variant clusters). By averaging the variant frequencies within each cluster for a given sample, we then analyzed the relative contribution of each cluster to the overall composition of the five lobes (Figure 5E-F). The degree of correlation between lobes (Figure 6B) is echoed by their degree of variant cluster sharing (lobe 1 shares cluster 4 with lobe 5, lobes 2 and 4 share variant cluster 8 with lobe 5, lobe 3 share clusters 7 and 8 with lobe 5) (Figure 5E-F). The presence of cluster 8 in four out of five lobes is especially notable, as it comprised a large percentage of the mutational burden in these four lobes (Figure 5E-F). Cluster 8 was defined by mutations in *Mll3* (also known as *Kmt2c*), *Setd2* and *Trp53* (Figure 5F). Variant-level analyses therefore recaptured the pairwise correlations identified on the sgRNA level, suggesting clonal mixture between individual liver lobes within a single mouse.

**Figure 5:**
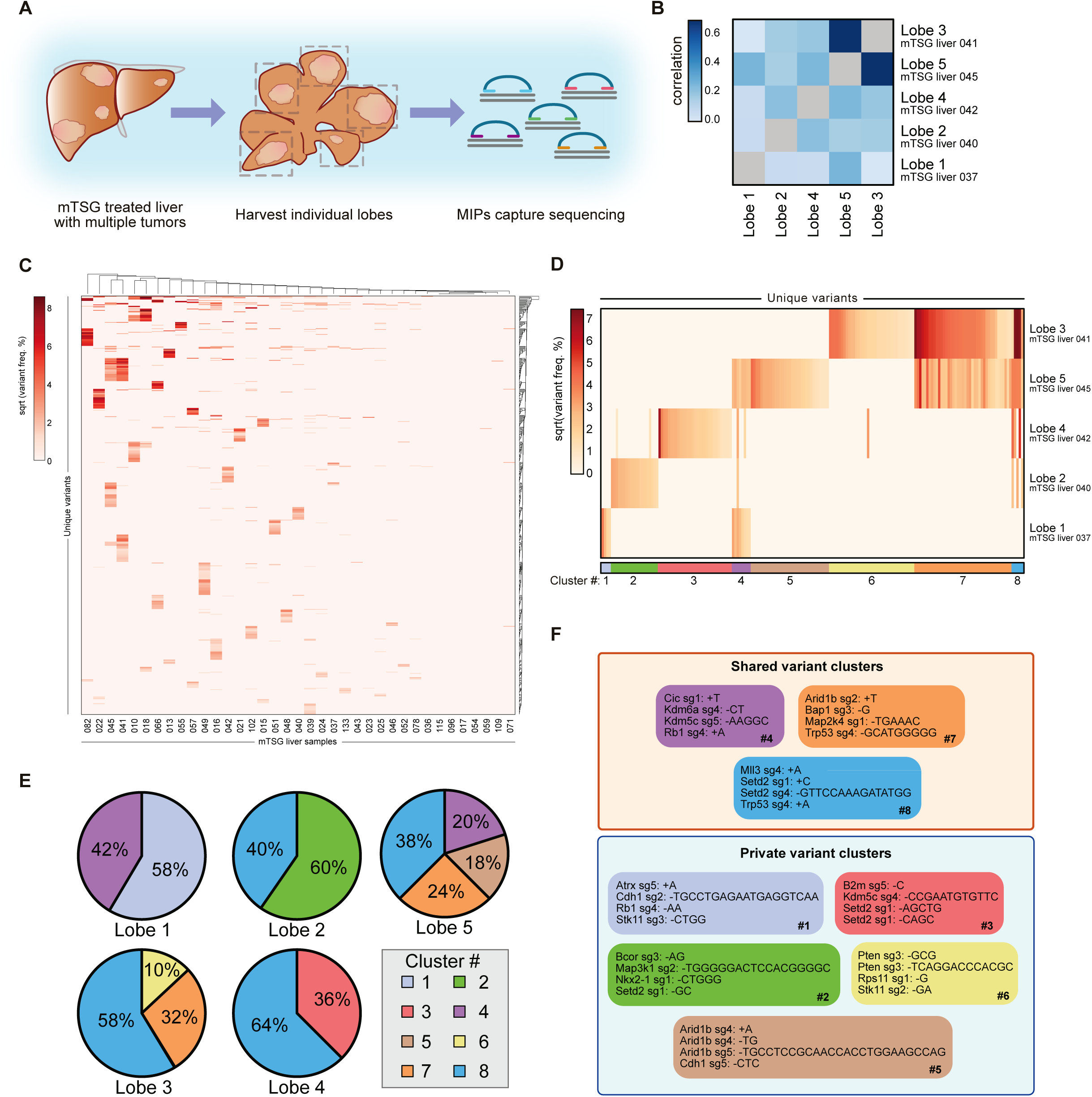
Systematic dissection of variant compositions across individual lobes within single animal. **A.** Schematic of the experimental workflow for analysis of multiple liver lobes (n = 5) from a single mTSG-treated mouse. **B.** Heatmap of Spearman’s rank correlation coefficients among five liver samples from a single mTSG-treated mouse, calculated on the basis of variant frequency for all unique variants present within the five samples. Notably, lobes 1-4 are all significantly correlated with lobe 5, with lobe 3 having the strongest correlation to lobe 5. **C.** Heatmap of all unique variants across all mTSG liver samples. Variant frequencies for all unique variants identified across mTSG liver samples, after square-root transformation for visual clarity. Rows denote unique variants, while columns denote different liver samples. Data was clustered using Euclidean distance and average linkage. 70.15% (416/593) of the variants were sample-specific, while 29.85% (177/593) variants were found across multiple samples. **D.** Heatmap of variant frequencies for each unique variant identified across the five individual liver lobes after square-root transformation. Rows correspond to different liver lobes, while columns denote unique variants. Eight clusters were identified based on binary mutation calls and are indicated on the bottom of the heatmap. **E.** Pie charts depicting the proportional contribution of each cluster to the five liver lobes. In order for a cluster to be considered, at least half of the variants within the cluster must be present in that particular sample. For each lobe, variant frequencies within a cluster were averaged and converted to relative proportions, as shown in the pie charts. The pie charts accurately recapture the correlation analysis in **b**, while additionally providing quantitative insight into the shared variants between the five liver lobes. **F.** Each box corresponds to one cluster, color-coded as in **c** and **d**, showing the top four variants in each cluster. On the basis of whether a variant cluster was present in multiple liver lobes, each box is also classified as either a private or a shared variant cluster. Clusters 1, 2, 3, 5 and 6 are largely unique to individual lobes (private variant clusters), while clusters 4, 7 and 8 are present in multiple lobes (shared variant clusters). Cluster 8 was found in four out of five lobes, and is characterized by mutations in *Mll3, Setd2* and *Trp53.*

We individually tested the functional roles of mutations in several of the top genes, in a *Trp53*-sensitized background. We generated an AAV vector for liver-specific CRISPR knockout that expressed Cre recombinase under a *TBG* promoter, together with a *Trp53-*targeting sgRNA cassette and an open (GeneX-targeting) sgRNA cassette (Figure S10A). The vector also contained a firefly luciferase gene (FLuc) co-cistronic with Cre under the *TBG* promoter for live imaging of tumorigenesis in mice. We cloned either a non-targeting control (NTC) sgRNA (thus only mutating *Trp53*), or a top candidate geneX-targeting sgRNA (GTS, thus mutating both *GeneX* and *Trp53*) into the 2^nd^ sgRNA expression cassette. After AAV packaging, we injected NTC + Trp53 or GTS + Trp53 AAVs into LSL-Cas9 mice (Figure 7A). We assessed the growth of potential liver tumors by monitoring their luciferase activities using a bioluminescent *in vivo* imaging system (IVIS) (Figure S10B and Figure S10D). Compared to mice treated with NTC AAVs (n = 8), sgRNAs targeting multiple candidates identified in the screen, including *Cic* (n = 4), *Pik3r1* (n = 7), *Pten* (n = 4), *Stk11* (n = 8), *Arid2* (n = 3), and *Kdm5c* (n = 3) had significantly stronger luciferase activity (two-sided unpaired t test, *p* < 0.05 for all groups, see Figure 7B-C), suggesting that knocking out these genes accelerated liver tumorigenesis at high penetrance in a *Trp53-*sensitized background. Double knockouts such as *Pik3r1 + Pten* (n = 3) and *Arid2 + Kdm5c* (n = 4) also had significantly stronger luciferase activity compared to NTC (two-sided unpaired *t* test, *p* < 0.001), but not significant compared to respective single knockouts (Figure S10B-C), suggesting that these genes are strong drivers alone but do not have synergistic effect with each other. Notably, the LSL-Cas9 mice injected with AAVs targeting *B2m + Kansl1*, one of the top co-occurring gene pairs identified in the screen (co-occurrence rate = 11/18 = 61%, *p* = 3.6* 10^-4^), showed significantly higher and increasing luminescence intensities compared with the mice injected with AAVs targeting NTC (two-sided unpaired t test, *p* < 0.01), *B2m* (*p* < 0.01) or *Kansl1* alone (*p* < 0.05), whereas mice with individual knockout of *B2m* or *Kansl1* alone did not show significantly stronger luminescence intensities than NTC controls (Figure S10B-D). These results suggested that combinatorial knockout of *B2m* and *Kansl1* had a synergistic effect in accelerating liver tumor development, whereas the single knockouts of *B2m* or *Kansl1* were not sufficient to induce liver tumorigenesis in a *Trp53*-sensitized background. In summary, the single and combinatorial AAVCRISPR knockout experiments validated the top ranked genes and co-occurring gene pairs in liver tumorigenesis. Our study collectively demonstrated a powerful strategy for quantitatively mapping functional landscapes and validating causality of TSGs and their synergistic relationships directly *in vivo*.

## Discussion

We developed a new technological approach, termed PASTAS, combining pooled AAV-CRISPR screens and targeted amplicon sequencing, which has broad applicability and several technological advantages over existing alternatives. First, AAVs are potent and safe carriers of transgenes compared to other types of viral vectors. Second, PASTAS directly mutagenizes the native cells in the endogenous organ and enables deep analysis of functional variants in an autochthonous manner, without relying on exogenous transplants. Third, compared to conventional sgRNA sequencing, MIP capture sequencing enabled direct, multiplexed analysis of the indels induced by Cas9 mutagenesis. By refocusing our analysis to the variant level, we systematically dissected the variant compositions across multiple liver lobes from a single mouse, uncovering evidence of clonal mixture between lobes. Whereas traditional sgRNA sequencing can only provide information about the relative abundances of each sgRNA, capture sequencing enables high-resolution analysis of individual indel variants for clonal analysis of tumor heterogeneity.

The present demonstration of the PASTAS method in LIHC shows that this approach can be capitalized to identify and validate causative genetic variants for massively parallel testing of the functional roles ^15,16,14,2,27,28^ in a quantitative and unbiased manner. This approach can potentially be extended to identify genetic factors with a significant impact on various cancer types and other human diseases. Our strategy for selecting genes to target in the mTSG library was based on pan-cancer TCGA datasets, with an initial aim of identifying genes that are more likely to function as tumor suppressors in a wide variety of tissues, and the overarching goal that the same AAVCRISPR mTSG library could potentially be used in other organs. In light of the previous success of *in vivo* transposon-based, shRNA or lentiviral screens in colon, liver and skin cancers ^55–61^ we anticipate that our approach can be readily expanded to other organ systems, potentially enabling the construction of a multi-organ functional variant mapping of tumor suppressors. Several *in vitro* gene activation screens using CRISPR have been described, opening the possibility for overexpression screens of proto-oncogenes ^61–65^. CRISPR has also recently been engineered to mutate specific DNA nucleotides ^64–66^. Adapting these new tools *in vivo* in conjunction with PASTAS would allow for high-throughput oncogene screens, thereby enabling functional variant mapping of oncogenes.

In summary, the Pooled AAV Screen with Targeted Amplicon Sequencing approach can be applied to any gene sets, any cancer types, or be focused down to the level of individual patients, personalized to the mutations present in each patient’s cancer, to empower therapeutic testing. All such studies can be performed in combination with many pre-clinical or co-clinical settings, providing brand new and powerful avenues for therapeutic discovery. Finally, with customized *in vivo* assay settings and readouts, the PASTAS approach can be applied to many types of *in vivo* genetic studies other than cancer.

## Acknowledgments

We thank all members in the Chen, Sharp, Zhang and Platt laboratories, as well as various colleagues in the Yale Department of Genetics, Systems Biology Institute, Yale Cancer Center and Stem Cell Center, Koch Institute and Broad Institute at MIT for assistance and/or discussion. We thank the Center for Genome Analysis, Center for Molecular Discovery, High Performance Computing Center, West Campus Analytical Chemistry Core and West Campus Imaging Core and Keck Biotechnology Resource Laboratory at Yale, as well as Swanson Biotechnology Center at MIT, for technical support.

## Contributions

SC and RJP planned, designed and initiated the experiments. GW and SC performed the majority of molecular, cellular, and animal work. RC and SC designed and performed the MIPS readout for functional cancer genomics and analyzed the data. LY, CDG, XD and MBD contributed to parts of the experiments. FZ, PAS, RJP and SC jointly supervised the work. SC and RC conceived and developed the functional genome variant mapping concept. RC prepared the first manuscript draft. RC, GW and SC wrote the paper with inputs from all authors.

## Methods

### Design, synthesis and cloning of the mTSG library

Pan-cancer mutation data from 15 cancer types were retrieved from The Cancer Genome Atlas (TCGA portal) via cBioPortal ^68,^^69^ and Synapse (https://www.synapse.org). Significantly mutated genes were calculated similarly to previously described methods ^14–16,38^. Known oncogenes were excluded and only known or predicted tumor suppressor genes (TSGs) were included. The top 50 TSGs were chosen, and their mouse homologs (mTSG) were retrieved from mouse genome informatics (MGI) (http://www.informatics.jax.org). A total of 49 mTSGs were found. A total of seven known housekeeping genes were chosen as internal controls. sgRNAs against these 56 genes were designed using a previously described method ^51,52^ with our custom scripts. Five sgRNAs were chosen for each gene, plus eight non-targeting controls (NTCs), making a total 288 sgRNAs in the mTSG library. NTCs do not target any predicted sites in the genome, thus were not included in subsequent MIPS analysis. Of note, two sgRNA pairs happened to be identical by design, namely Rpl22_sg4/sg5, and Cdkn2a_sg2/sg5. These sgRNAs were treated as the same in subsequent analyses.

### Design, cloning of AAV-CRISPR vectors and mTSG sgRNA library cloning

AAV-CRISPR vectors were designed to express Cre recombinase for the induction of Cas9 expression using constitutive or conditional promoters when delivered to LSL-Cas9 mice ^9^. Two sgRNA cassettes were built in these vectors, one encoding an sgRNA targeting *Trp53*, with the other being an open sgRNA cassette (double SapI sites for sgRNA cloning). The vector was generated by gBlock gene fragment synthesis (IDT) followed by Gibson assembly (NEB). The mTSG library were generated by oligo synthesis, pooled and cloned into the double SapI sites of the AAV-CRISPR vectors. Library cloning was done at over 100x coverage to ensure proper representation. Representation of plasmid libraries was readout by barcoded Illumina sequencing as described previously ^53^ with customized primers.

### AAV-mTSG viral library production

The AAV-CRISPR plasmid vector (AAV-vector) and library (AAV-mTSG) were subjected to AAV9 production and chemical purification. Briefly, HEK 293FT cells (ThermoFisher) were transiently transfected with AAV-vector or AAV-mTSG, AAV9 serotype plasmid and pDF6 using polyethyleneimine (PEI). Each replicate consists of five 80% confluent HEK 293FT cells in 15-cm tissue culture dishes or T-175 flasks (Corning). Multiple replicates were pooled to enhance production yield. Approximately 72 h post transfection, cells were dislodged and transferred to a conical tube in sterile PBS. 1/10 volume of pure chloroform was added and the mixture was incubated at 37°C and vigorously shaken for 1 h. NaCl was added to a final concentration of 1 M and the mixture was shaken until dissolved and then pelleted at 20,000 g at 4°C for 15 min. The aqueous layer was discarded while the chloroform layer was transferred to another tube. PEG8000 was added to 10% (w/v) and shaken until dissolved. The mixture was incubated at 4°C for 1 h and then spun at 20,000 g at 4° C for 15 min. The supernatant was discarded and the pellet was resuspended in DPBS plus MgCl_2_ and treated with Benzonase (Sigma) and incubated at 37°C for 30 min. Chloroform (1:1 volume) was then added, shaken, and spun down at 12,000 g at 4°C for 15 min. The aqueous layer was isolated and passed through a 100 kDa MWCO (Millipore). The concentrated solution was washed with PBS and the filtration process was repeated. Genomic copy number (GC) of AAV was titrated by real-time quantitative PCR using custom Taqman assays (ThermoFisher) targeted to Cre.

### Animal work statements

All animal work was performed under the guidelines of Yale University Institutional Animal Care and Use Committee (IACUC) and Massachusetts Institute of Technology Committee for Animal Care (CAC), with approved protocols (Chen-2015-20068 and Sharp-0914-091-17), and were consistent with the Guide for Care and Use of Laboratory Animals, National Research Council, 1996 (institutional animal welfare assurance no. A-3125- 01).

### Intravenous (i.v.) virus injection for liver transduction

Conditional LSL-Cas9 knock-in mice were bred in a mixed 129/C57BL/6 background. Mixed gender (randomized males and females) 8-14 week old mice were used in experiments. Mice were maintained and bred in standard individualized cages with maximum of five mice per cage, with regular room temperature (65-75°F, or 18- 23°C), 40-60% humidity and a 12h:12h light cycle. To intravenously inject AAVs, mice were restrained in rodent restrainer (Braintree Scientific), their tails were dilated using a heat lamp or warm water, sterilized by 70% ethanol, and 200 μL of concentrated AAV (~1e10 GC/μL, 2e12 GC per mouse) was injected into the tail vein of each mouse. 100% of the mice survived the procedure. Animals that failed injections (< 70% of total volume injected into tail vein after multiple attempts) were excluded from the study. No specific methods were implemented to choose sample sizes.

### MRI

MRI imaging was performed using standard imaging protocol with MRI machines (Varian 7T/310/ASR-whole mouse MRI system, or Bruker 9.4T horizontal small animal systems). Briefly, animals were anesthetized using isoflurane, and positioned in the imaging bed with a nosecone providing constant isoflurane. A total of 20 - 30 frontal views were acquired for each mouse using a custom setting: echo time (TE) = 20, repetition time (TR) = 2000, slicing = 1.0 mm. Raw image stacks were processed using Osirix or Slicer tools ^70^. Rendering and quantification were performed using Slicer (www.slicer.org). Tumor size was calculated with the following formula: Volume (mm^3^) = 1/6 * 3.14 * length (mm) * height (mm) * depth (mm). Statistical significance was assessed by non-parametric Mann-Whitney test, as samples numbers and sample distributions varied across treatment conditions.

### Survival analysis

We observed that LSL-Cas9 mice receiving AAV-mTSG intravenous injections rapidly deteriorate in their body condition scores (due to tumor development in most cases). Mice with body condition score (BSC) < 2 were euthanized and the euthanasia date was recorded as the last survival date. Occasionally mice bearing tumors died unexpectedly early, and the date of death was recorded as the last survival date. Cohorts of mice intravenously injected with PBS, AAV-vector or AAV-mTSG virus were monitored for their survival. Survival analysis was analyzed by standard Kaplan – Meier method, using the *survival* and *survminer* R packages. Differences among the three treatment groups were assessed by log-rank test. Of note, several AAV-vector or PBS injected mice were sacrificed at time points earlier than the last day of survival analysis (at times when a certain AAV-mTSG mice were found dead or euthanized due to poor body conditions), to provide time-matched histology, even though those mice presented with good body condition (BSC >= 4). Mice euthanized early in a healthy state were excluded from calculation of survival percentages.

### Mouse organ dissection, fluorescent imaging and histology

Mice were sacrificed by carbon dioxide asphyxiation or deep anesthesia with isoflurane followed by cervical dislocation. Mouse livers and other organs were manually dissected and examined under a fluorescent stereoscope (Zeiss, Olympus or Leica). Brightfield and/or GFP fluorescent images were taken for the dissected organs and overlaid using ImageJ ^71^. Organs were then fixed in 4% formaldehyde or 10% formalin for 48 to 96 hours, embedded in paraffin, sectioned at 6 μm and stained with hematoxylin and eosin (H&E) for pathology. For tumor size quantification, H&E slides were scanned using an Aperio digital slidescanner (Leica). Tumors were manually outlined as region-of-interest (ROI), and subsequently quantified using ImageScope (Leica). Statistical significance was assessed by Welch’s t-test, given the unequal sample numbers and variances for each treatment condition.

### Mouse tissue collection for molecular biology

Mouse livers and various other organs (supplemental tables) were dissected and collected manually. For molecular biology, tissues were flash frozen with liquid nitrogen, ground in 24 Well Polyethylene Vials with metal beads in a GenoGrinder machine (OPS diagnostics). Homogenized tissues were used for DNA/RNA/protein extractions using standard molecular biology protocols.

### Genomic DNA extraction from cells and mouse tissues

For genomic DNA extraction, 50-200 mg of frozen ground tissue were resuspended in 6 mL of Lysis Buffer (50 mM Tris, 50 mM EDTA, 1% SDS, pH 8) in a 15 mL conical tube, and 30 μL of 20 mg/mL Proteinase K (Qiagen) were added to the tissue/cell sample and incubated at 55 °C overnight. The next day, 30 μL of 10 mg/mL RNAse A (Qiagen) was added to the lysed sample, which was then inverted 25 times and incubated at 37 °C for 30 min. Samples were cooled on ice before the addition of 2 mL of pre-chilled 7.5 M ammonium acetate (Sigma) to precipitate proteins. The samples were vortexed at high speed for 20 s and then centrifuged at ≥ 4,000 *g* for 10 min. Then, a tight pellet was visible in each tube and the supernatant was carefully decanted into a new 15 ml conical tube. Then 6 ml 100% isopropanol was added to the tube, inverted 50 times and centrifuged at ≥ 4,000 *g* for 10 min. Genomic DNA was visible as a small white pellet in each tube. The supernatant was discarded, 6 ml of freshly prepared 70% ethanol was added, the tube was inverted 10 times, and then centrifuged at ≥ 4,000 *g* for 5 min. The supernatant was discarded by pouring; the tube was briefly spun, and remaining ethanol was removed using a P200 pipette. After air-drying for 10-30 min, the DNA changed appearance from a milky white pellet to slightly translucent. Then, ~500 μL of ddH2O was added, the tube was incubated at 65 °C for 1 h and at room temperature overnight to fully resuspend the DNA. The next day, the genomic DNA samples were vortexed briefly. The concentration of genomic DNA was measured using a Nanodrop (Thermo Scientific).

### Molecular Inversion Probe (MIP) design and synthesis

MIPs were designed according to previously published protocols ^72,73^. Briefly, the 70 bp flanking the predicted cut site of each sgRNA of all 278 unique sgRNA were chosen as targeting regions, and the bed file with these coordinates was used as an input. Since *Trp53* sg4 targets a similar region as the *p53* sgRNA within the base vector, the same MIP was used to sequence both of these loci.

These coordinates contained overlapping regions which were subsequently merged into 173 unique regions. Each probe contains an extension probe sequence, a ligation probe sequence, and a 7 bp degenerate barcode (NNNNNNN) for PCR duplicate removal. A total of 266 MIP probes were designed covering a total amplicon of 42,478 bp. MIP target size stats: min = 155 bp, max = 190 bp, mean = 159.7 bp, median = 156.0 bp (supplemental table). Each of the mTSG-MIPs were synthesized using standard oligo synthesis (IDT), normalized and pooled.

### MIP capture sequencing

150 ng of genomic DNA sample from each mouse organ was used as input. MIP capture sequencing was done according to previously published protocols ^72,73^ with some slight modifications. The multiplexed library was then quality controlled using qPCR, and subjected to high-throughput sequencing using the Hiseq-2500 or Hiseq-4000 platforms (Illumina) at Yale Center for Genome Analysis. 280/281 (99.6%) of targeted sgRNAs were captured for all samples from this experiment, with the missing one being Arid1a sg5.

### Illumina sequencing data pre-processing

FASTQ reads were mapped to the mm10 genome using the bwa mem function in BWA v0.7.13 ^74^. Bam files were merged, sorted, and indexed using bamtools v2.4.0 ^75^ and samtools v1.3 ^76^.

### Variant calling

For each sample, indel variants were called using samtools and VarScan v2.3.9 ^77^. Specifically, we used samtools mpileup (–d 1000000000 –B –q 10), and piped the output to VarScan pileup2indel (--min-coverage 1 -- min-reads2 1 --min-var-freq 0.001 --p-value 0.05). To link each indel to the sgRNA that most likely caused the mutation, we took the center position of each indel and mapped it to the closest sgRNA cut site.

### Calling significantly cutting sgRNAs and significantly mutated genes

We further filtered all detected indels by requiring that each indel must overlap the ± 3 basepair flank of the closest sgRNA cut site, as Cas9-induced double-strand breaks are expected to occur within a narrow window of the predicted cut site. To exclude any possible germline mutations, we also removed any sgRNAs with indels present in more than half of the control samples with greater than 5% variant frequency. In particular, high variant frequencies were observed across all samples at the *Rps19* sg5 cut site, suggesting these were germline variants; thus, we excluded *Rps19* sg5 from all analyses.

To determine significantly mutated sgRNA sites in each liver sample, we used a false-discovery approach based on the PBS and vector control samples. For each sgRNA, we first took the highest % variant read frequency across all control liver samples; in order for a mutation to be called in an mTSG sample, the % variant read frequency had to exceed the control sample cutoff. However, since the base vector contained a *Trp53* sgRNA (p53 sg8) whose cut site was only 1 bp away from the target site of *Trp53* sg4 (from mTSG library), we only considered PBS samples when calculating the false-discovery cutoff for *Trp53* sg4. Finally, as we were most interested in identifying the dominant clones in each sample, we further set a 5% variant frequency cutoff on top of the false-discovery cutoff. These criteria gave us a binary table (i.e. not significantly mutated vs. significantly mutated) detailing each sgRNA and whether its target site was significantly mutated in each sample. To convert significantly mutated sgRNA sites into significantly mutated genes, we simply collapsed the binary sgRNA scores by gene, such that if any of the five sgRNAs for a gene were found to be significantly cutting, the entire gene would be called as significantly mutated.

### Coding frame analysis

For coding frame and exonic/intronic analysis, we only considered the indels that were associated with an sgRNA which had been considered significantly mutated in that particular sample. This final set of significant indels was converted to avinput format and subsequently annotated using ANNOVAR v. 2016Feb01, using default settings ^78^.

### Cooccurrence and correlation analysis

Cooccurrence analysis was performed by first generating a double-mutant count table for each pairwise combination of genes in the mTSG library. Statistical significance of the cooccurrence was assessed by two-sided hypergeometric test. To calculate cooccurrence rates, we defined the “intersection” as the number of double-mutant samples, and the “union” as the number of samples with a mutation in either (or both) of the two genes, and then divided the intersection by the union. For correlation analysis, we first collapsed the table of variant frequencies to the gene level (in other words, summing the variant frequencies for all five of the targeting sgRNAs for each gene). Using these summed variant frequency values, we calculated the Spearman or Pearson correlation between all gene pairs, across each mTSG sample. Statistical significance of the correlation was determined by converting the correlation coefficient to a t-statistic, and then using the t-distribution to find the associated probability. For both cooccurrence and correlation analyses, *p*-values were adjusted for multiple hypothesis testing by the Benjamini-Hochberg method to obtain *q*-values.

### Unique variant analysis

Instead of first collapsing variant calls to the sgRNA level as above, unique variants and their associated mutant frequencies were compiled across all sequenced samples. To be considered present in a given sample, a particular variant must have a mutant frequency >= 1%. Heatmaps of the unique variant landscape were created in R using the *NMF* package, with average linkage and Euclidean distance. We also performed a focused analysis on the unique variant landscape within a single mouse. For the correlation heatmap, we used Spearman rank correlation to assess the pairwise correlation between different liver lobes. Clusters of variants were defined on the basis of binary mutation calls (i.e. whether a given variant is present or not within each sample). To determine the proportional contribution of each cluster, for each sample, we only included the clusters in which at least half of the variants in the cluster are present in that sample. We then took the average mutant frequency across the variants within each cluster, and used these values to determine the relative contribution of each cluster to the overall sample. To identify the top four variants in each cluster, we ranked all the variants by the average variant frequency across all lobes in which the variant cluster was considered present.

### Direct in vivo validation of drivers or combinations

Liver-specific AAV-CRISPR vectors were designed to co-cistronically express firefly luciferase (FLuc) and Cre recombinase for induction of Cas9 expression under a *TBG* promoter when delivered to LSL-Cas9 mice (Plasmids available at Addgene). Two sgRNA cassettes were built in these vectors, one encoding an sgRNA targeting *Trp53*, with the other being an open sgRNA cassette (double SapI sites for GeneX targeting sgRNA cloning). The vector was generated by gBlock gene fragment synthesis (IDT) followed by Gibson assembly (NEB). Each specific sgRNA targeting a driver gene was cloned separately into this vector. AAV9 virus was produced and qPCR-titrated as described above. 1e11 total viral particles were introduced by intravenous injection into LSL-Cas9 mice. For combinations of two AAVs, 5e10 viral particles were used from each AAV to generate equal titer mixtures and injected. Four to six mice were injected per group. Starting one month after injection, mice were imaged by IVIS each month. Briefly, mice were anesthetized by intraperitoneal (I.P.) injection of ketamine (100 mg/kg) and xylazine (10 mg/kg), and imaged for *in vivo* tumor growth using an IVIS machine (PerkinElmer) with 150 mg/kg body weight Firefly D-Luciferin potassium salt injected I.P. Relative luciferase activity was quantified using LivingImage software (PerkinElmer).

### Blinding statement

Investigators were blinded for histology scoring and MIPS, but not blinded for dissection, MRI or survival analysis.

### Code availability

All custom scripts used to process and analyze the data will be available after publication.

### Accessions

Genomic sequencing data are being deposited in NCBI SRA under a pending accession number.

CRISPR reagents (plasmids and libraries) are being deposited to Addgene to share with the academic community.

## Supplementary Figure Legends

**Figure S1:**
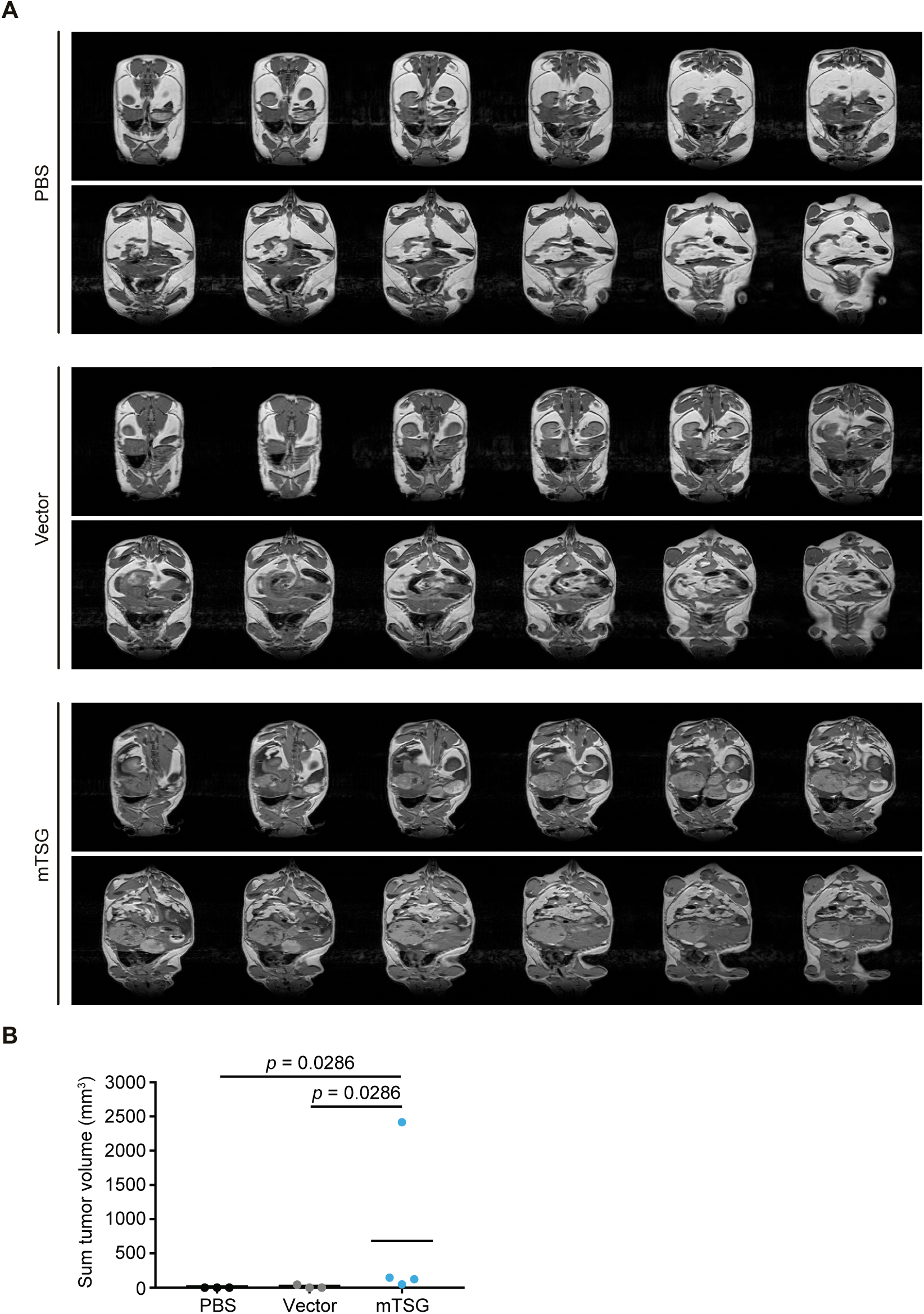
Representative full-spectrum MRI series of livers from PBS, vector and mTSG-treated mice. **A.** Full-spectrum MRI slices from representative PBS, vector, and mTSG-treated mice. **B.** Dot plot of the sum tumor volume per mouse (in mm^3^) in mice treated with PBS (black, n = 3), vector (gray, n = 3), or mTSG library (blue, n = 4). mTSG-treated mice had significantly higher tumor burdens than PBS (one-sided Mann-Whitney test, *p* = 0.0286) or vector-treated animals (*p* = 0.0286). **(related to Figure 1)**

**Figure S2:**
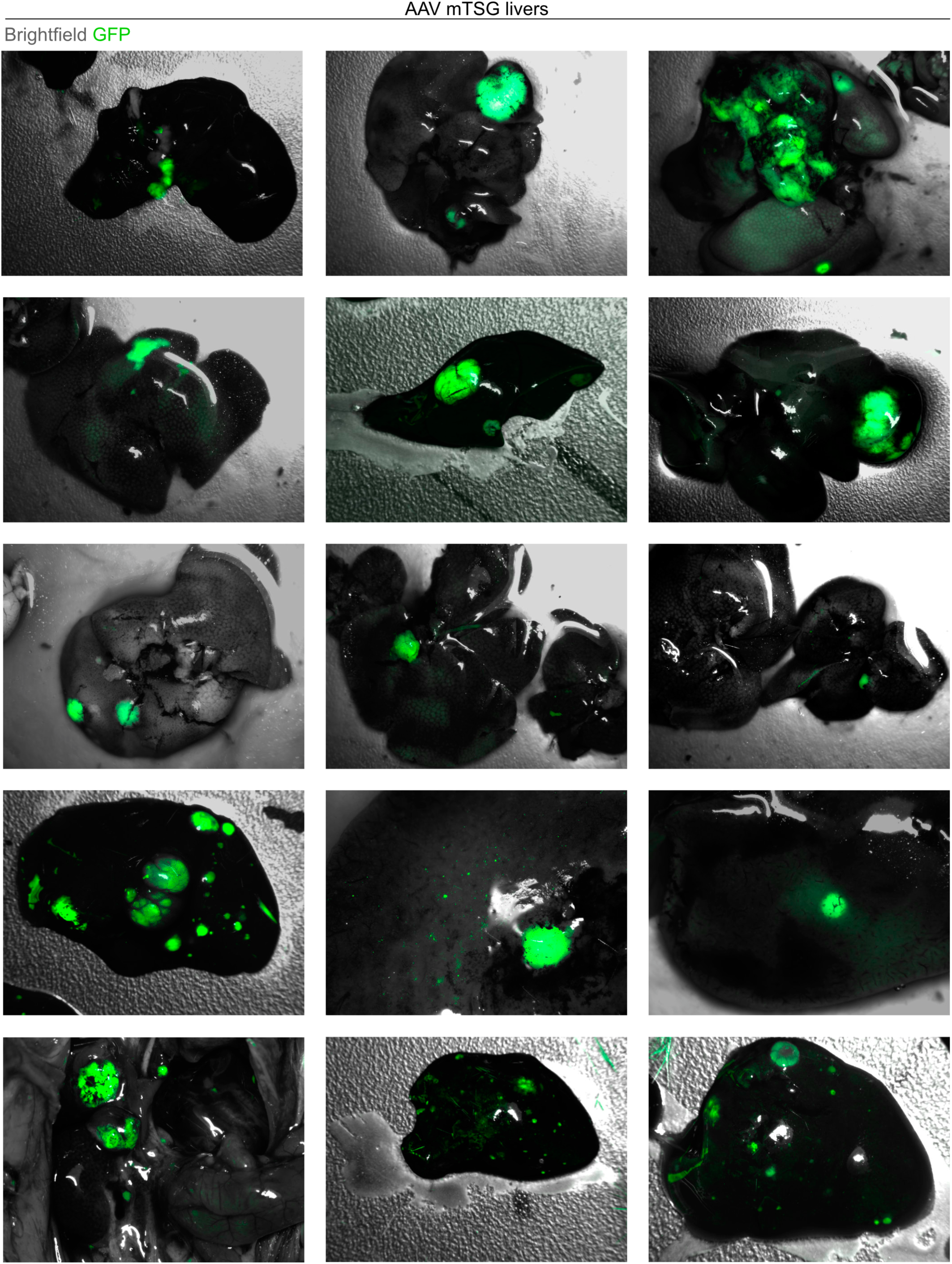
Additional brightfield images of mTSG-treated livers with GFP overlay. Additional brightfield images with GFP fluorescence overlay (green) of livers from 15 mTSG-treated mice at the time of sacrifice.

**Figure S3:**
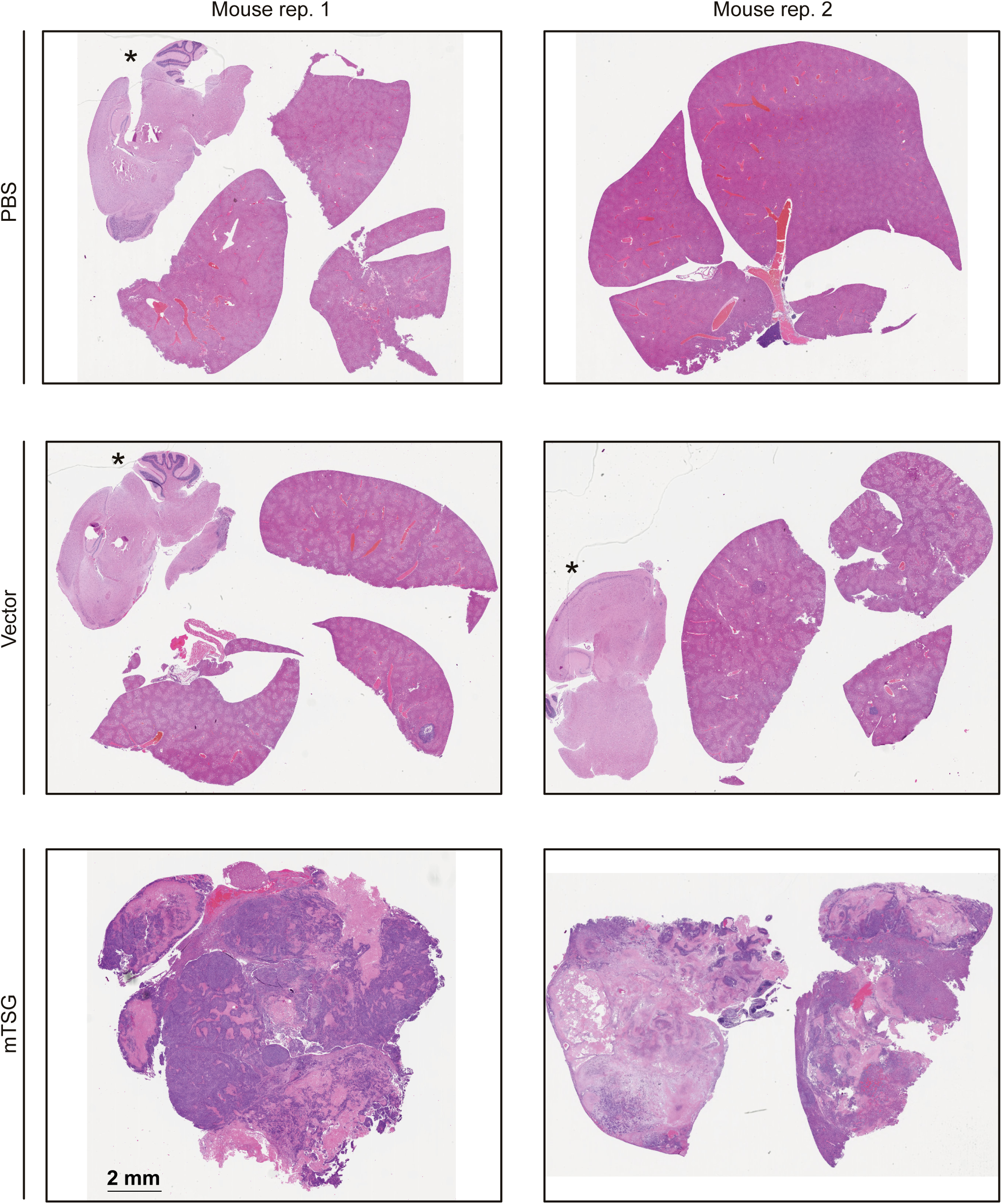
Representative full slide scanning images of mouse liver sections in PBS, vector and mTSG treatment groups. Full slide scans of liver sections from PBS, vector and mTSG-treated mice. Two representative mice from each group are shown. Some brain sections are also present in the same scanned field, noted with asterisks. PBS samples did not have any detectable nodules, while vector-treated samples occasionally had developed small nodules. In contrast, mTSG-treated samples were replete with tumors. Scale bar is 2 mm.

**Figure S4:**
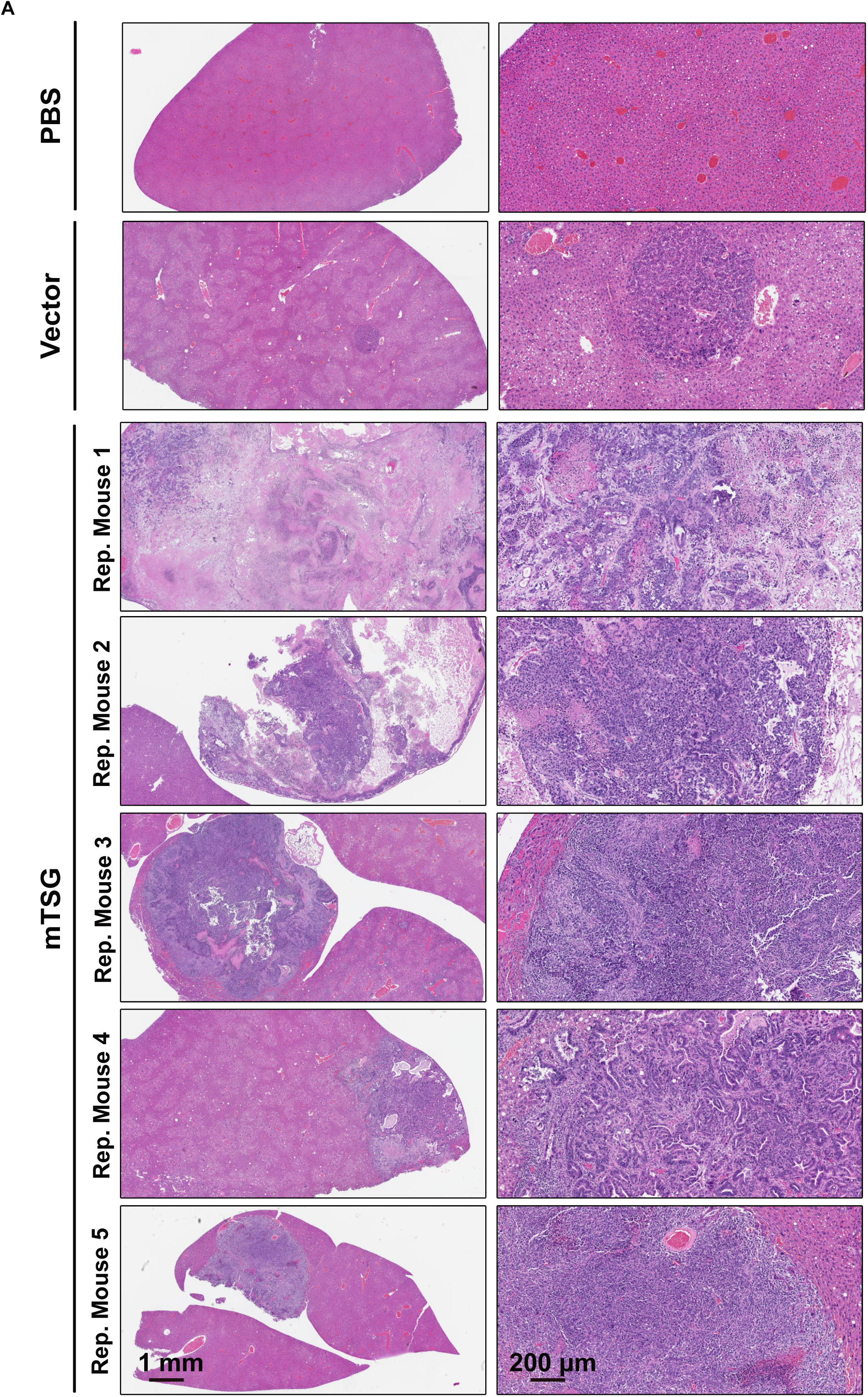
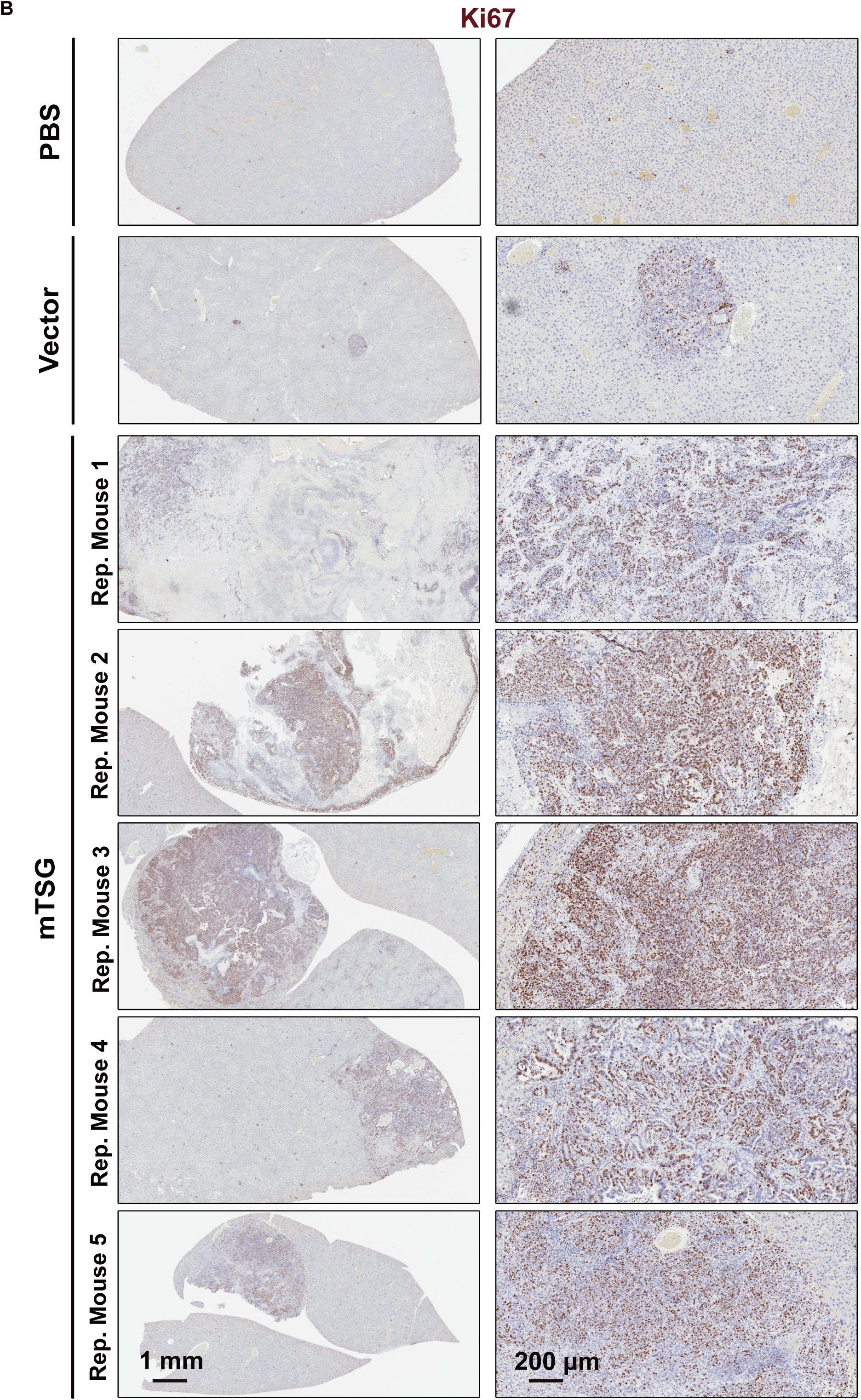
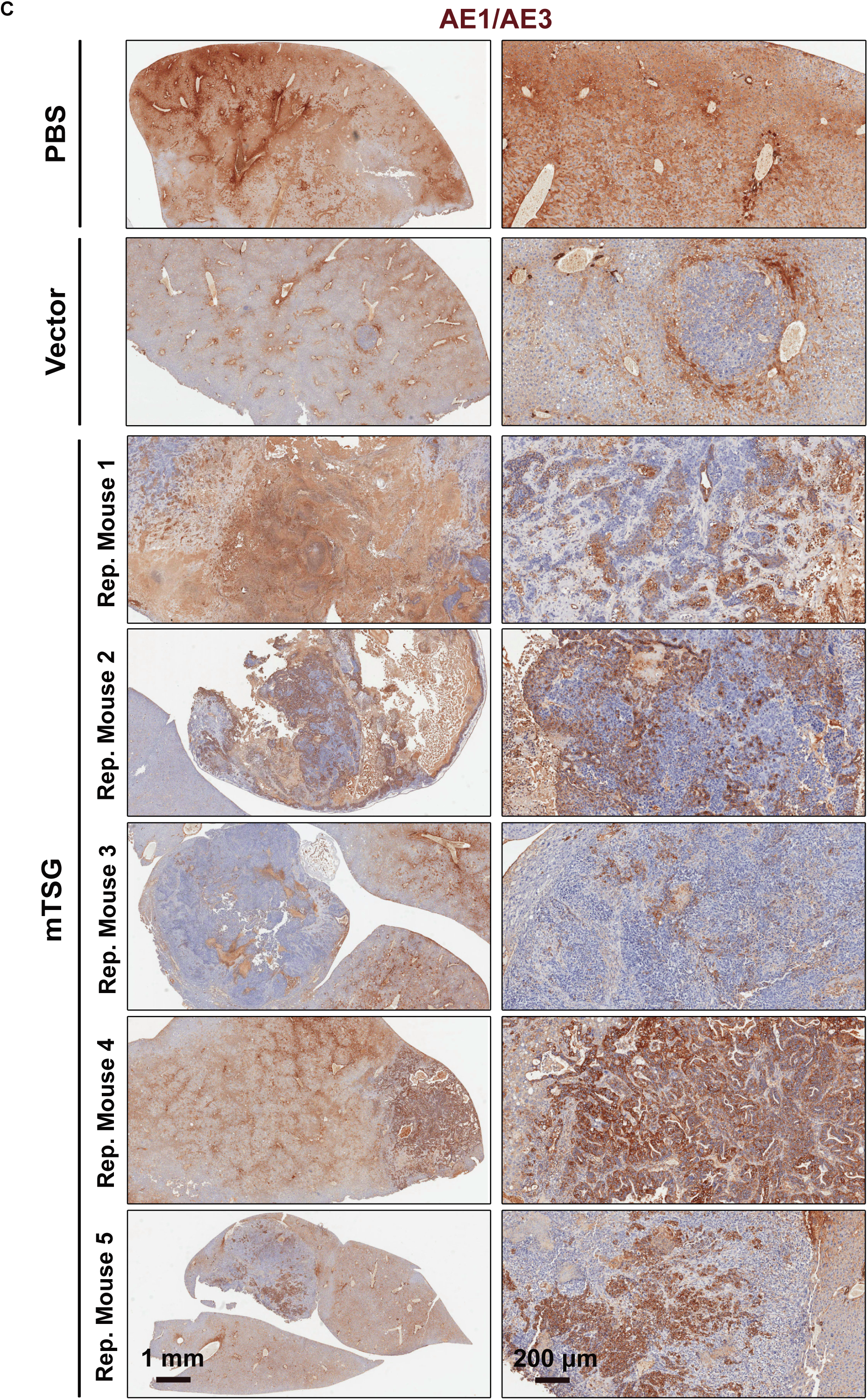
Representative histology and immunohistochemistry images of mouse liver sections in PBS, vector and mTSG groups. **A.** Representative liver sections from PBS, vector, and mTSG-treated mice with hematoxylin and eosin staining. The vector sample and mTSG replicate 4 pictured here are from the same mice shown in Figure 2D. Scale bar is 1 mm for low magnification images, 200 μm for high magnification images. **B.** Representative liver sections from PBS, vector, and mTSG-treated mice with Ki67 staining. Sections correspond to the same mice shown in Fig. S4A. Scale bar is 1 mm for low magnification images, 200 μm for high magnification images. **C.** Representative liver sections from PBS, vector, and mTSG-treated mice with pan-cytokeratin AE1/AE3 staining. Sections correspond to the same mice shown in Fig. S4A. Scale bar is 1 mm for low magnification images, 200 μm for high magnification images.

**Figure S5:**
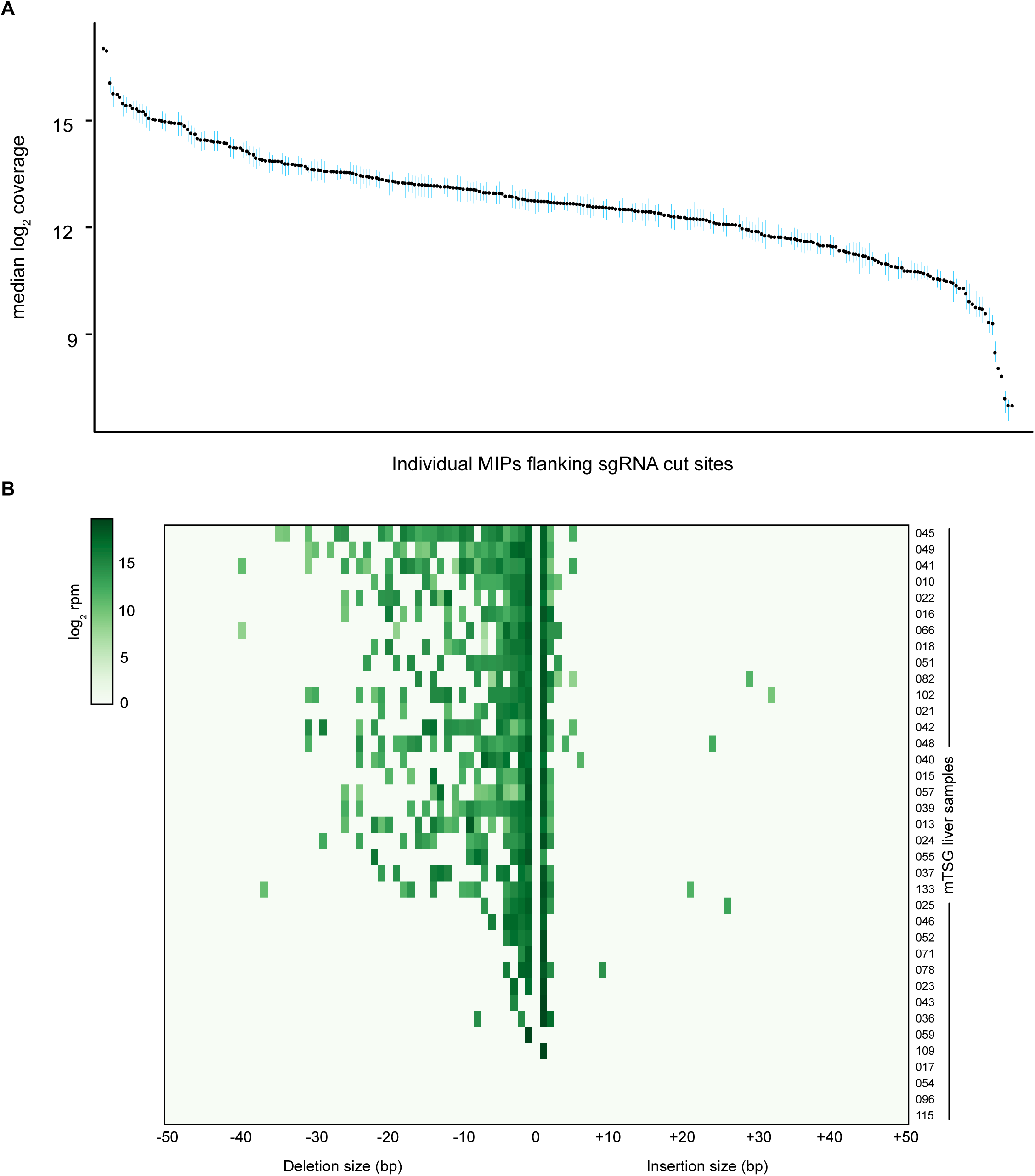
MIPs capture sequencing statistics and indel size distribution of mTSG livers. **A.** Plot of median log_2_ sequencing coverage across all sequenced samples in amplicons targeted by the 266 MIPs (black dots). MIPs were designed to amplify the genomic regions flanking the predicted cut sites of each sgRNA. 95% confidence intervals for the median are depicted with blue lines. Median read depth across all MIPs approximated a lognormal distribution, indicating relatively even capture of the target loci. **B.** Heatmap detailing indel size distribution and abundance across all significantly mutated sgRNA sites from mTSG-treated liver samples. Positive indel sizes denote insertions, while negative indel sizes indicate deletions. Depicted values are in terms of total log2 normalized reads per million (rpm) for each sample. Most variant reads are deletions (80.8%) compared to insertions (19.2%).

**Figure S6:**
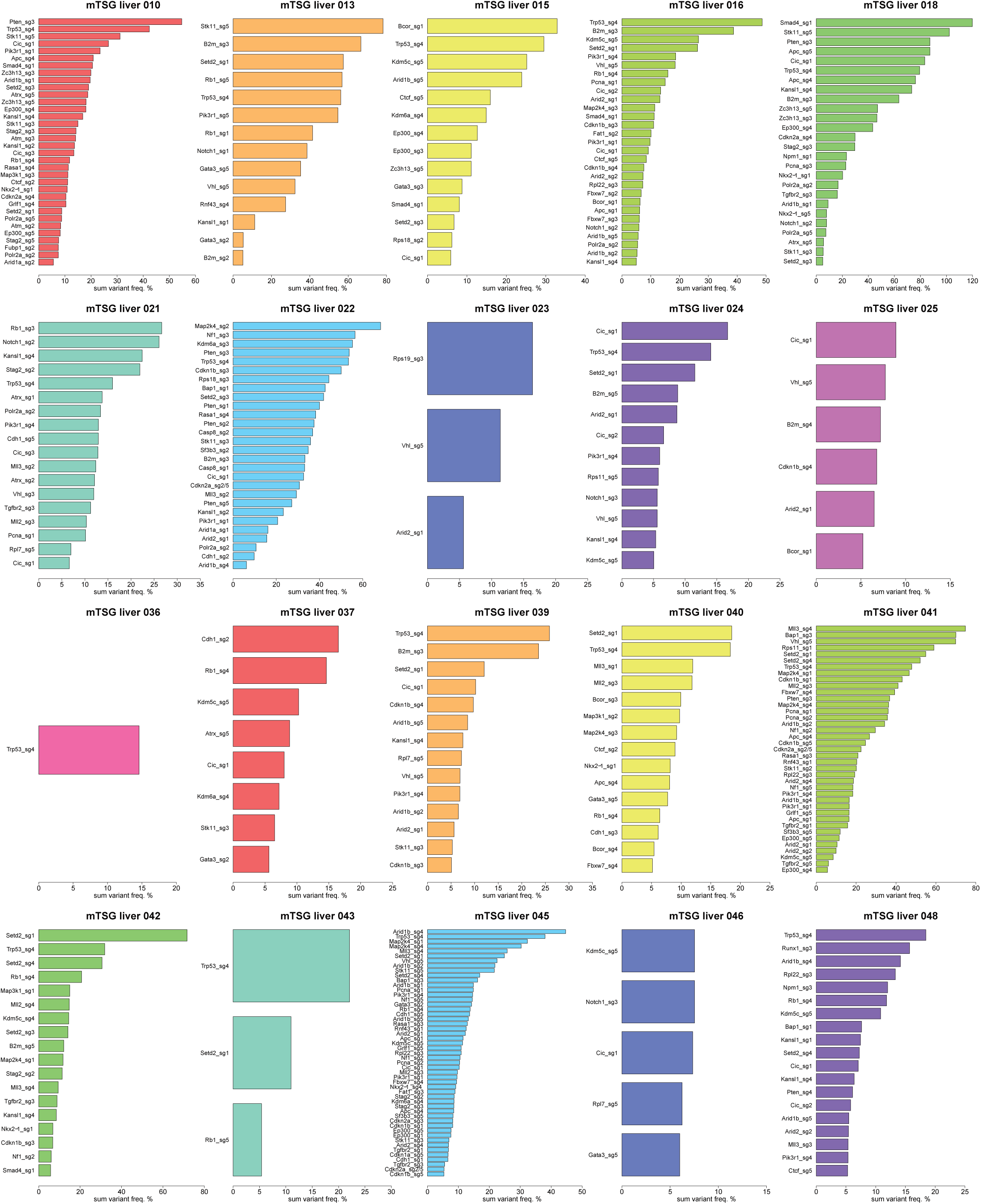
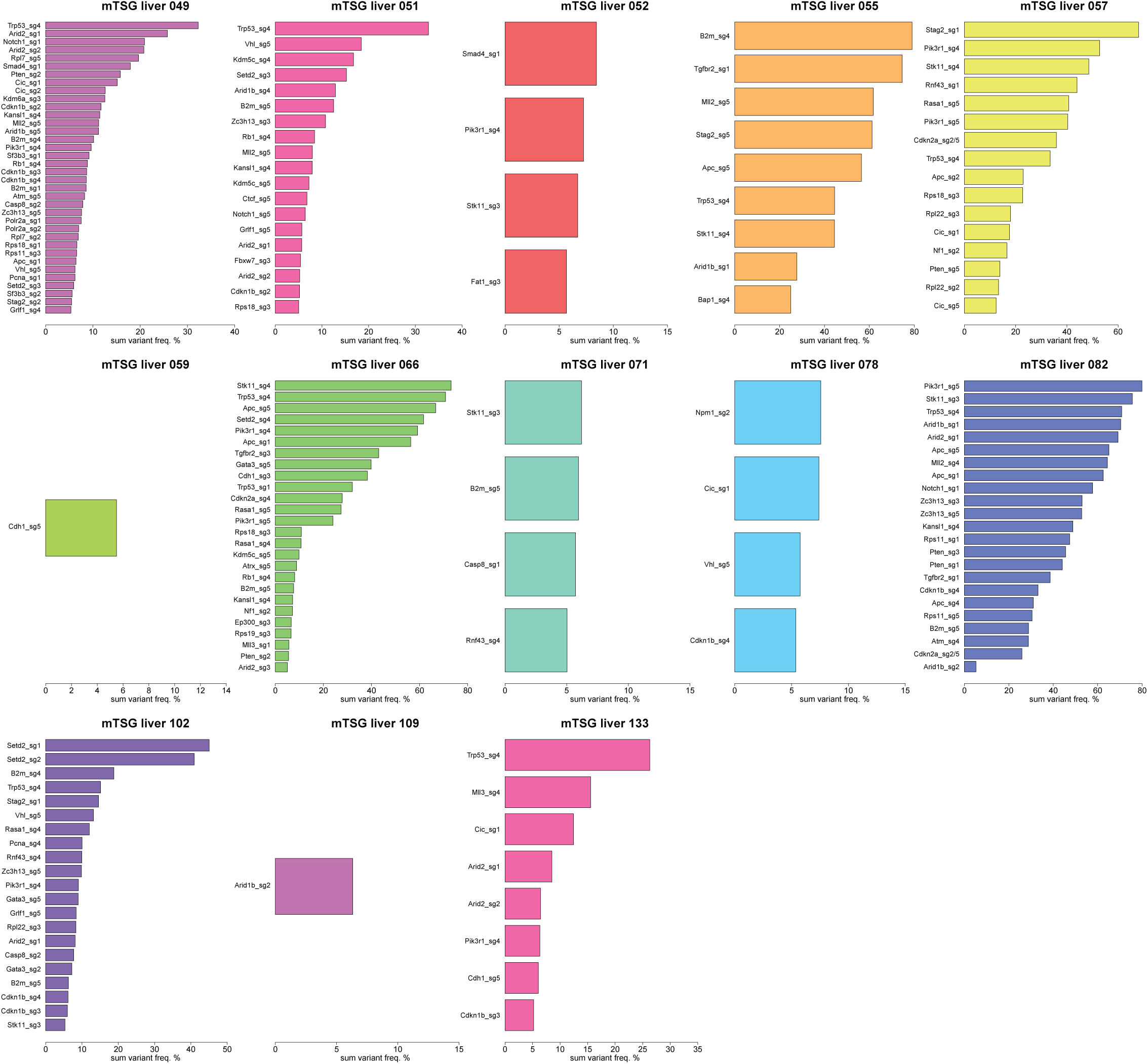
Significantly mutated sgRNA sites across all liver samples from mice treated with AAV-mTSG library. Waterfall plots of significantly mutated sgRNA sites across all 33 mTSG-treated liver samples, sorted by sum variant frequency. Four samples (mTSG liver 17, mTSG liver 54, mTSG liver 96 and mTSG liver 115) are not shown, as these samples were not found to have any significantly mutated sgRNA sites per our stringent variant calling strategy. The extensive mutational heterogeneity amongst the liver samples is suggestive of strong positive selective forces acting on diverse loss-of-function mutations induced by the mTSG library.

**Figure S7:**
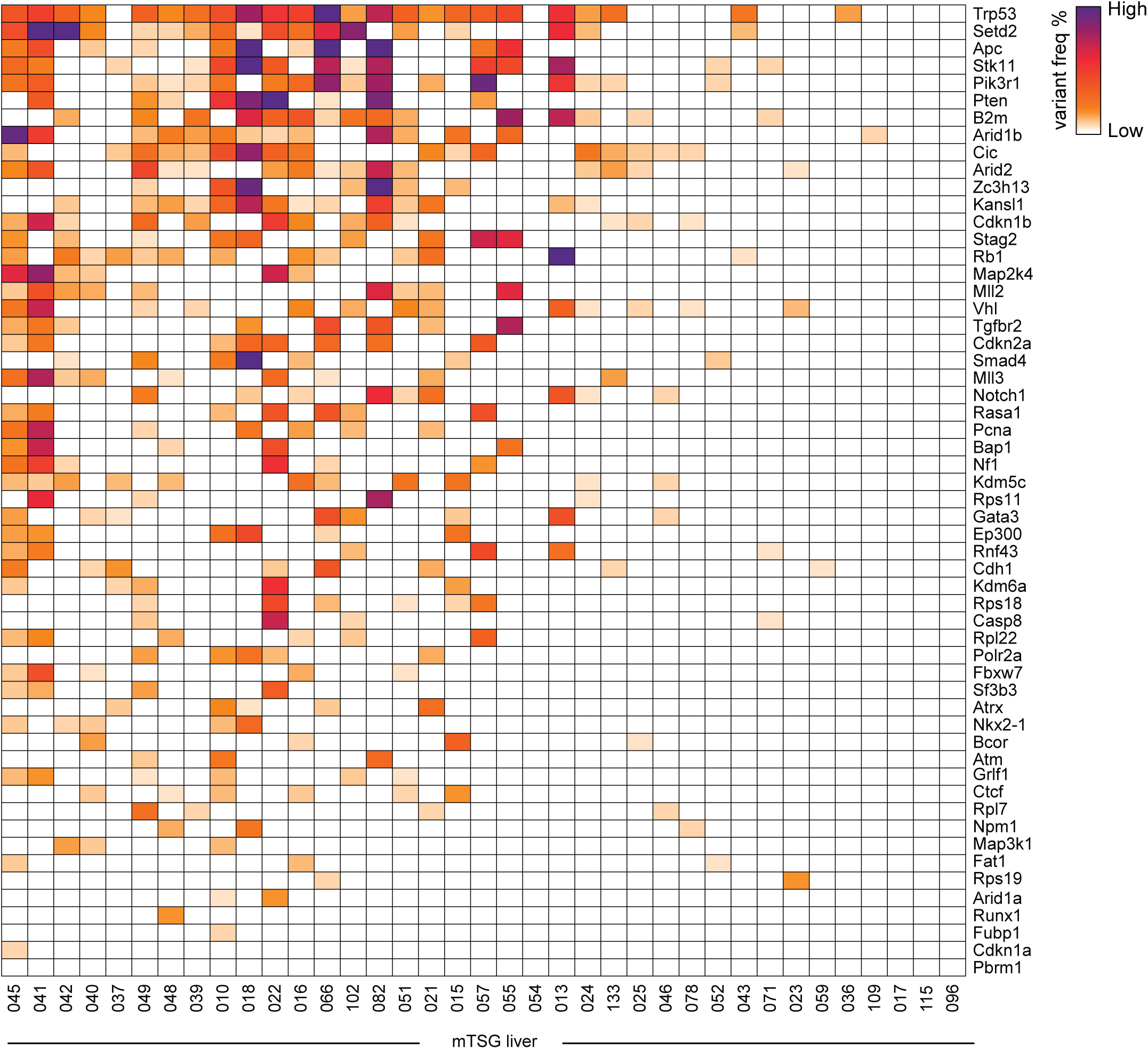
Heatmap of gene level sum variant frequency across all mTSG liver samples. Heatmap depicting sum variant frequencies for the 56 genes represented in the library, across all mTSG liver samples. Genes are ordered according to average sum variant frequency (top to bottom row).

**Figure S8:**
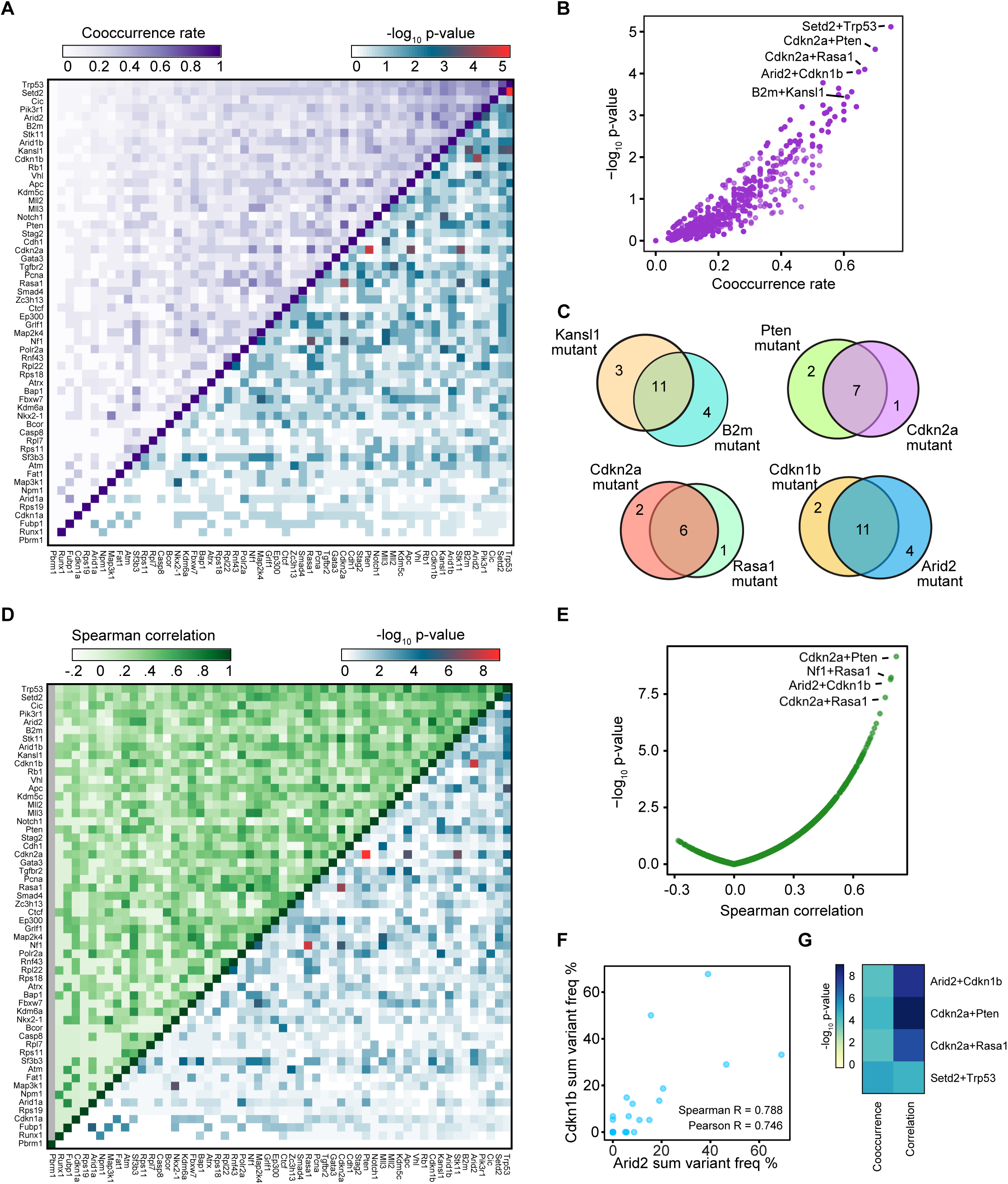
Co-mutation analysis of synergistic combinations of driver mutations. **A.** Upper-left triangle: heatmap of the cooccurrence rates for each gene pair. To calculate cooccurrence rates, the “intersection” is defined as the number of double-mutant samples, and the “union” as the number of samples with a mutation in either of the two genes. The cooccurrence rate was then calculated as the intersection divided by the union. Lower-right triangle: heatmap of -log_10_ p-values by hypergeometric test to evaluate whether specific pairs of genes are statistically significantly co-mutated. **B.** Scatterplot of the cooccurrence rates for each gene pair, plotted against -log_10_ p-values by hypergeometric test. Highly co-occurring pairs include *Cdkn2a* + *Pten* (co-occurrence rate = 7/10 = 70%; hypergeometric test, *p* = 2.63 * 10^-5^), *Cdkn2a* + *Rasa1* (6/9 = 67%; *p* = 7.96 * 10^-5^), *Arid2* + *Cdkn1b* (11/17 = 65%; *p* = 9.13 * 10^-5^) and *Kansl1 + B2m* (11/18 = 61%; *p* = 3.6 * 10^-4^). **C.** Venn diagrams showing the strong co-occurrence of mutations in *Setd2* + *Trp53* (top left), *Cdkn2a + Pten* (top right), *Cdkn2a + Rasa1* (bottom left), and *Arid2 + Cdkn1b* (bottom right). Numbers shown correspond to the number of mTSG-treated liver samples with a given mutation profile. **D.** Upper-left triangle: heatmap of the pairwise Spearman correlation of sum % variant frequency for each gene, summed across sgRNAs. Lower-right triangle: heatmap of -log_10_ p-values by t-distribution to evaluate the statistical significance of the pairwise correlations. **E.** Scatterplot of pairwise Spearman correlations plotted against -log_10_ p-values. The top four correlated pairs were *Cdkn2a* + *Pten* (Spearman R = 0.817, *p* = 6.97* 10^-10^), *Nf1* + *Rasa1* (R = 0.791, *p* = 5.86 * 10^-9^), *Arid2* + *Cdkn1 b* (R = 0.788, *p* = 7.16 * 10^-9^), and *Cdkn2a* + *Rasa1* (R = 0.761, *p* = 4.45 * 10^-8^). **F.** Scatterplot comparing sum level % variant frequency for *Arid2* vs. *Cdkn1 b* across all mTSG-treated liver samples. Spearman and Pearson correlation coefficients are noted on the plot (Spearman R = 0.788; Pearson R = 0.746). **G.** Heatmap of the p-values associated with the top mutation pairs that were found to be statistically significant (Benjamini-Hochberg adjusted *p <* 0.05) in both cooccurrence (left) and correlation (right) analyses.

**Figure S9:**
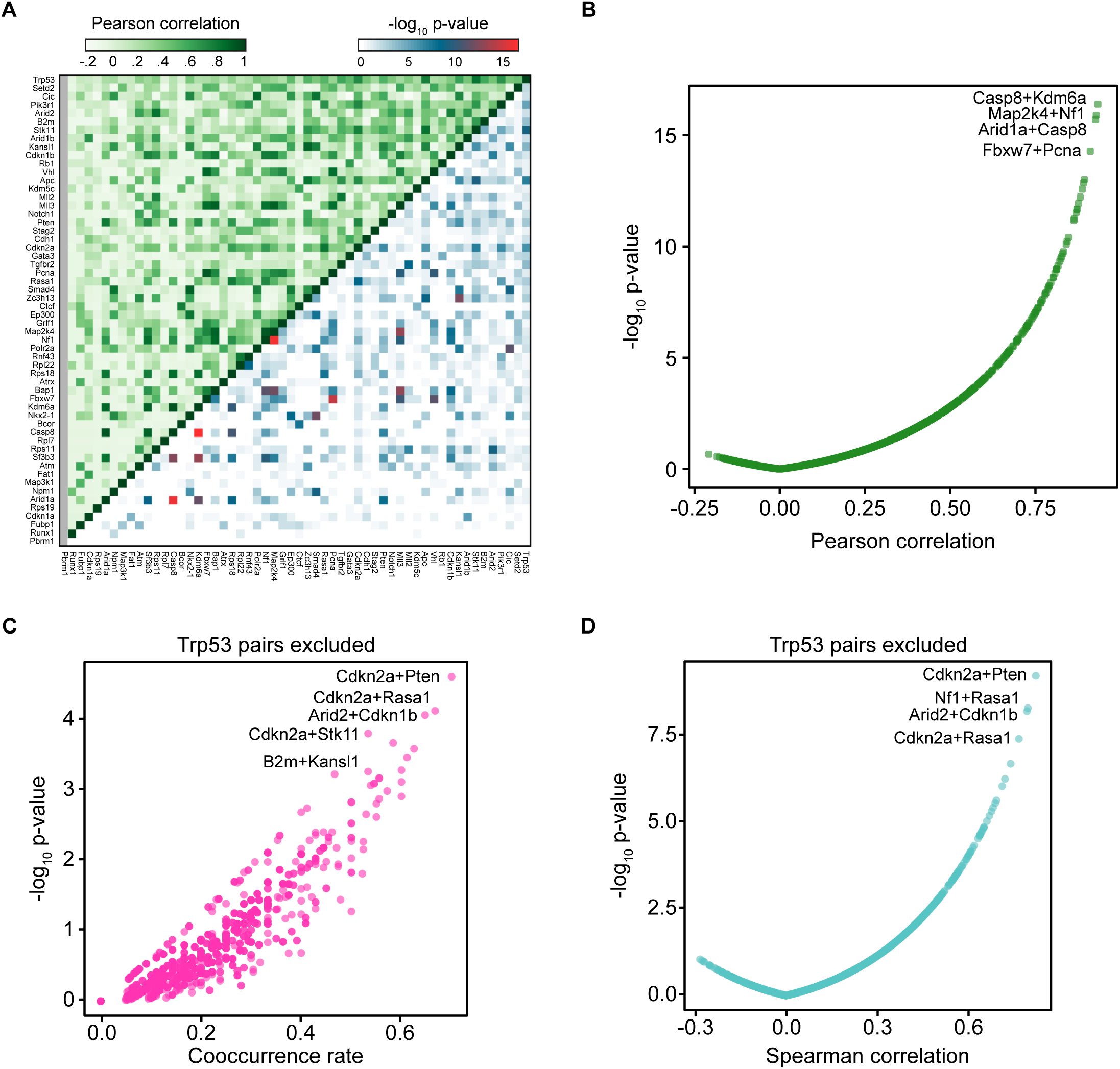
Additional co-mutation analysis. **A.** Upper-left triangle: heatmap of the pairwise Pearson correlation of sum % variant frequency for each gene, averaged across sgRNAs. Lower-right triangle: heatmap of -log_10_ p-values by t-distribution to evaluate the statistical significance of the pairwise correlations. **B.** Scatterplot of the Pearson correlation for each gene pair, plotted against -log_10_ p-values. **C.** Scatterplot of the cooccurrence rates for each gene pair, excluding all pairs involving *Trp53*, plotted against - log_10_ p-values by hypergeometric test. **D.** Scatterplot of the Spearman correlations for each gene pair, excluding all pairs involving *Trp53*, plotted against - log_10_ p-values.

**Figure S10:**
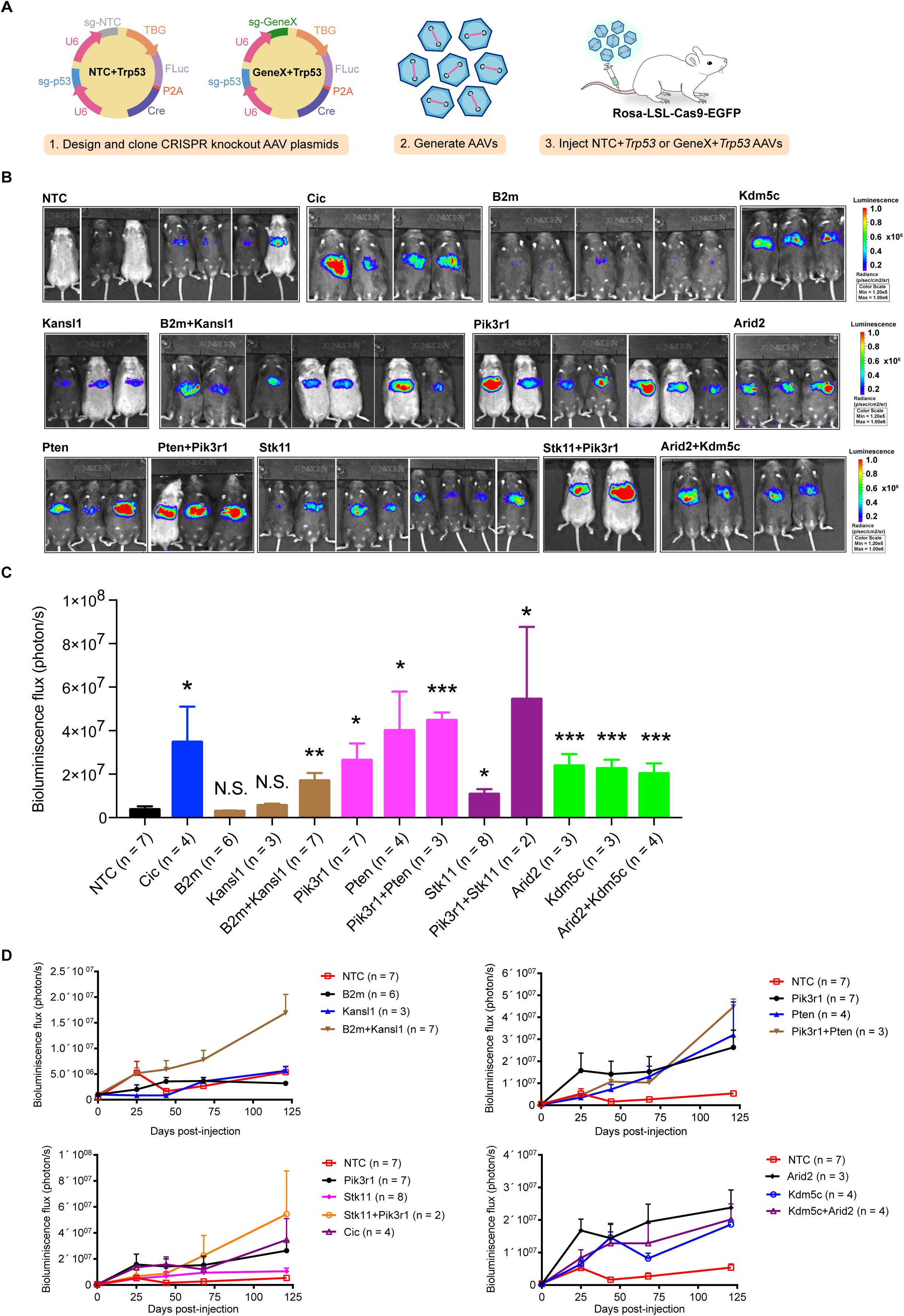
Single or combinatorial AAV-CRISPR knockout of TSGs in driving liver tumorigenesis. **A.** Schematics of the design and cloning of liver-specific AAV-CRISPR vectors to functionally study target genes for their potential roles as independent and synergistic drivers of liver tumor in immunocompetent mice. The AAVCRISPR plasmids contain two U6 promoter-driving sgRNA expression cassettes, with the 1^st^ sgRNA targeting *Trp53,* and another one either as a non-targeting sgRNA (NTC + *Trp53*) or a geneX-targeting sgRNA (GeneX + *Trp53*). The plasmids also contained a liver-specific TBG promoter driving a co-cistronic expression cassette of firefly luciferase (FLuc) and Cre recombinase. AAVs were generated with these plasmids and injected intravenously into LSL-Cas9 mice. **B.** Representative bioluminescence images of LSL-Cas9 mice injected with AAV9 that contains liver-specific TBG promoter-driving Cre and CRISPR dual-sgRNAs expression cassettes. Undetectable or weak luciferase activity was detected in NTC + *Trp53* AAV treated mice (n = 8) at 121 days post-injection, whereas persistent and robust luciferase activity was detected in the mice that were injected with the top scoring genes (GeneX + *Trp53*) or the highly co-mutated gene pairs from the screen. **C.** Quantification of bioluminescence intensities of AAV-CRISPR injected LSL-Cas9 mice at 121 days post-injection are shown in units of photons/sec/cm^2^/sr (Data represented as mean ± SEM). The mice that were injected with AAVs targeting the top screened genes or the highly correlated gene pairs had robust luciferase activity after 121 days of injection, indicating the role of these TSGs in accelerating development of tumors compared to NTC controls (two-sided unpaired *t* test, N.S. *p* > 0.05, * *p* < 0.05, ** *p* < 0.01, *** *p* < 0.001). In comparison to NTC (n = 7), *Cic* (n = 4, *p* = 0.018), *Pik3r1* (n = 7, *p* = 0.015), *Pten* (n = 4, *p* = 0.011), *Stk11* (n = 8, *p* = 0.03), *Arid2* (n = 3, *p* = 0.001) and *Kdm5c* (n = 3, *p* = 0.0005) knockout had significantly higher bioluminescence intensities. Double knockout of *Pik3r1+Pten* (n = 3) had significantly stronger luciferase activity compared to NTC (two-sided unpaired t test, *p* < 0.0001), but was not significantly different from knocking out *Pik3r1* or *Pten* alone (two-sided unpaired t test, N.S.). Double knockout of *Pik3r1+Stk11* (n = 2) had significantly stronger luciferase activity compared to NTC (two-sided unpaired *t* test, *p* = 0.01), but was not significantly different from knocking out *Pik3r1* or *Stk11* alone (two-sided unpaired *t* test, N.S.). In contrast, double knockout of *B2m+Kansl1* led to significantly higher luminescence intensities compared to NTC (two-sided unpaired *t* test, *p* = 0.005), *B2m* alone (*p* = 0.001) and *Kansl1* alone (*p* = 0.02). **D.** Longitudinal IVIS live imaging of bioluminescence intensities of LSL-Cas9 mice injected with liver-specific AAVs containing either NTCs or sgRNAs targeting a single gene or a combination of two genes, including the following pairs: *B2m + Kansl1*, *Pik3r1* + *Pten, Pik3r1* + *Stk11* and *Arid2 + Kdm5c.* Data are shown as mean ± SEM in units of photons/sec/cm^2^/sr.

## Supplementary Tables

Table S1. DNA sequences of sgRNA spacers in mTSG library.

Table S2. Raw read counts of mTSG plasmid library.

Table S3. Tumor volume data as measured by MRI.

Table S4. Survival data for PBS, vector, or mTSG-treated animals.

Table S5. Tumor area data as measured by tissue histology.

Table S6. Sequence information and annotation for all MIPs used in the study.

Table S7. Metadata for all of the 133 sequenced samples.

Table S8. MIPs capture sequencing coverage statistics across all predicted cutting sites of sgRNAs in AAV mTSG library.

Table S9. Raw indel variant calls of all samples with targeted capture sequencing before filtering.

Table S10. sgRNA level sum indel frequency table for all samples with targeted capture sequencing.

Table S11. sgRNA level binary SMS calls in livers from mice treated with AAV mTSG library.

Table S12. Gene level binary SMG calls in livers from mice treated with AAV mTSG library.

Table S13. Cooccurrence analysis of SMG pairs in livers from mice treated with AAV mTSG library.

Table S14. Correlation analysis of gene level sum indel frequency in livers from mice treated with AAV mTSG library.

Table S15. Mutant frequencies for all unique variants present across all mTSG liver samples.

Table S16. Mutant frequencies for all unique variants present in 5 individual liver lobes from a single mouse.

